# Histone chaperone HIRA, Promyelocytic Leukemia (PML) protein and p62/SQSTM1 coordinate to regulate inflammation during cell senescence

**DOI:** 10.1101/2023.06.24.546372

**Authors:** Nirmalya Dasgupta, Xue Lei, Christina Huan Shi, Rouven Arnold, Marcos G. Teneche, Karl N. Miller, Adarsh Rajesh, Andrew Davis, Valesca Anschau, Alexandre R. Campos, Rebecca Gilson, Aaron Havas, Shanshan Yin, Zong Ming Chua, Tianhui Liu, Jessica Proulx, Michael Alcaraz, Mohammed Iqbal Rather, Josue Baeza, David C. Schultz, Kevin Y. Yip, Shelley L. Berger, Peter D. Adams

## Abstract

Cellular senescence, a stress-induced stable proliferation arrest associated with an inflammatory Senescence-Associated Secretory Phenotype (SASP), is a cause of aging. In senescent cells, Cytoplasmic Chromatin Fragments (CCFs) activate SASP via the anti-viral cGAS/STING pathway. PML protein organizes PML nuclear bodies (NBs), also involved in senescence and anti-viral immunity. The HIRA histone H3.3 chaperone localizes to PML NBs in senescent cells. Here, we show that HIRA and PML are essential for SASP expression, tightly linked to HIRA’s localization to PML NBs. Inactivation of HIRA does not directly block expression of NF-κB target genes. Instead, an H3.3-independent HIRA function activates SASP through a CCF-cGAS-STING-TBK1-NF-κB pathway. HIRA physically interacts with p62/SQSTM1, an autophagy regulator and negative SASP regulator. HIRA and p62 co-localize in PML NBs, linked to their antagonistic regulation of SASP, with PML NBs controlling their spatial configuration. These results outline a role for HIRA and PML in regulation of SASP.

## Introduction

Senescence is a phenomenon characterized by stable cell-cycle arrest, which prevents the proliferation of damaged cells^1^. Senescence not only causes proliferation arrest but also involves epigenetic and metabolic reprogramming, as well as a pro-inflammatory secretome called the senescence-associated secretory phenotype (SASP)^2^. Senescence is considered a biomarker of aging and a key contributor to cell and tissue aging, leading to age-related diseases such as renal dysfunction, type-2 diabetes, Idiopathic pulmonary fibrosis (IPF), metabolic dysfunction-associated fatty liver disease, osteoarthritis, and declining immune function^3,4^. SASP is recognized as a contributing factor to the development of senescence-related and age-related diseases, tissue damage, and degeneration. This is due to its ability to act as a source of chronic inflammation or “inflammaging” in aged tissue^5^. The contribution of factors to SASP is dependent on the cellular context, although some key effectors and regulatory mechanisms are commonly shared. The transcription factor Nuclear Factor κB (NF-κB) functions as a “master regulator” of the SASP^6^. Additionally, CCAAT/enhancer–binding protein beta (C/EBPß) and GATA4 have also been identified as important transcriptional regulators of the SASP^3,7,8^. The SASP is regulated by various signaling molecules, including pathways such as p38 MAP-kinase, mTOR, JAK-STAT, and Notch, along with epigenetic regulators like G9a, GLP, macroH2A, HMGB2, MLL1, BRD4, and H2A.J^3,7,8^. SASP can also be regulated by DNA damage^9^. We have observed a phenomenon in senescent cells whereby damaged DNA is expelled from the nucleus as cytoplasmic chromatin fragments (CCF)^10^. Other DNAs in the cytoplasm of senescent cells include mitochondrial DNA^11^ and reverse transcribed retrotransposons^12^. These cytoplasmic DNAs trigger the activation of SASP through the cytosolic DNA-sensing cGAS-STING-NF-κB pathway^11-15^. Given that SASP is regulated at various levels, gaining a comprehensive understanding of its regulation is crucial. This understanding can pave the way for potential therapeutic approaches aimed at modulating SASP, which could promote healthy aging and alleviate the burden of senescence-associated diseases and degeneration.

Emerging evidence highlights the critical role of chromatin in integrating the SASP^7^. Previously we found that the chromatin in senescent cells is maintained in a state of dynamic equilibrium, and the HIRA chaperone complex plays a critical role in regulating the chromatin landscape of these cells^16^. The HIRA chaperone complex consists of HIRA, UBN1, and CABIN1, which work together with the histone-binding protein ASF1a to incorporate non-canonical H3.3 into chromatin in a manner that is independent of DNA replication^16^. The deposition of H3.3 through the HIRA pathway has been firmly linked to genes that are actively being transcribed^17^. Furthermore, it plays a crucial role in either activating or maintaining gene expression patterns in the long term^18^. When cells undergo senescence, their chromatin can become condensed, resulting in the formation of senescence-associated heterochromatin foci (SAHF)^19^. SAHF exhibit a distinct architecture comprising a core of constitutive heterochromatin encircled by a ring of facultative heterochromatin, with active chromatin located outside, and pericentromeric/telomeric regions positioned at the periphery^20^. This suggests a process of spatial chromatin refolding within SAHF^20^. Our observations have shown that HIRA plays a crucial role in the formation of SAHF ^21,22^ and that the overexpression of HIRA is sufficient to induce SAHF formation^21^. It was proposed that promyelocytic leukemia nuclear bodies (PML NBs) serve as a molecular “staging ground” where HIRA complexes are assembled or modified before being translocated to chromatin and contributing to the formation of SAHF^21,22^. While we know that HIRA is linked to cellular senescence, the exact function of HIRA in senescence is still not fully understood.

Prior to the onset of other senescence markers, such as cell cycle exit, SAHF, a large flat morphology, and SA β-Gal activity, HIRA enters within the subnuclear organelle PML NBs during senescence induction, and this relocation is essential for SAHF formation^21,22^. A typical senescent nucleus contains 10-30 PML NBs, which are spherical structures of 0.2-1 μm in diameter^21,23^. PML NBs play a versatile role in regulating a wide range of cellular activities, such as stress response, senescence, apoptosis, DNA repair, anti-viral immunity, and stem cell renewal^24^. The PML tumor-suppressor protein acts as the primary scaffold of PML NBs, crucial for maintaining the structural integrity of these bodies^25,26^. PML interacts with numerous other partner proteins, referred to as ’client’ proteins, to form the NBs^26^. However, the clients are dispensable for PML NBs and diffuse more rapidly than the ’scaffold’ PML protein^26^. PML can self-assemble through the binding of its conserved SUMO Interaction Motif (SIM) element to SUMOs (small ubiquitin-like modifiers) conjugated at multiple sites within the protein^26^. The SUMO-SIM interactions are crucial for recruiting numerous client proteins to PML NBs, including transcription factors, DNA repair proteins, enzymes involved in post-translational protein modifications^23,26^. An intriguing characteristic among these partner client proteins is their ability to undergo SUMOylation in PML NBs^23^. The level of PML protein increases in senescent cells^27,28^, and has been reported to play a key role in senescence induction by repressing the expression of E2F target genes^29^. In senescent cells, PML NBs are frequently found at the periphery of persistent DNA damage foci known as “DNA segments with chromatin alterations reinforcing senescence” (DNA-SCARS)^9^. The integrity of these DNA-SCARS contributes to the regulation of SASP^9^. Despite the presence of substantial evidence supporting the role of PML in senescence phenotypes, its precise function remains unclear and elusive.

Given the co-localization of HIRA and PML in senescent cells, in this study we investigated their concerted involvement in senescence and made several noteworthy findings. Firstly, we found that HIRA and PML are essential for the expression of senescence-associated secretory phenotype (SASP) in senescent cells, but their role is not linked to growth arrest. Our experiments revealed that the localization of HIRA to PML NBs is critical for the expression of SASP. Additionally, we made an intriguing discovery of a novel non-canonical function of HIRA, independent of its histone deposition activity, in regulating SASP expression.

## Results

### HIRA and PML are required for SASP expression but not for proliferation arrest

To begin to investigate the functions of HIRA and PML in senescence, we induced senescence in IMR-90 cells using etoposide and carefully monitored the accumulation of HIRA within PML NBs. During senescence, as previously shown by us and others^21,22,27,28^, PML NBs were observed to become larger and more numerous, followed by the later accumulation of HIRA in PML NBs (Figure S1A-D). When we disrupted the weak hydrophobic interactions in PML NBs of senescent cells using 1,6-hexanediol, we found that PML exhibited high resistance to dissolution in senescence (Figure S2A), consistent with previous observations during viral infection^30^. Therefore, PML is regarded as a scaffold protein in PML NBs in senescent cells^26^. In contrast, HIRA is more labile within PML NBs, as evidenced by the immediate dissolution of HIRA foci within one minute (Figure S2A). Surprising, other partner proteins in PML NBs, including SP100 and HP1IZ foci, are also stable and can be localized in PML NBs even after 15 minutes of 1,6-hexanediol treatment (Figure S2B,C). Investigating the association of PML NBs with DNA-SCARS by Super-Resolution confocal microscopy, we observed PML is not only at the periphery of the DNA-SCARS, rather it is frequently wrapped by DNA-SCARS (Figure S3A-C). A peak in PML staining intensity corresponds to a trough in γH2AX. HIRA was found to be much more closely colocalized with PML than was γH2AX (as shown in Figure S1A-D) and was also wrapped by DNA-SCARS (Figure S3D-F). DNA-SCARS are further distinguished by an increased presence of p53-binding protein 1 (53BP1)^9^. Similar to PML^9^, HIRA does not fully colocalize with 53BP1 (Figure S3G).

To investigate the role of HIRA and PML in senescence, we induced senescence after knocking down HIRA and PML using lentivirus-encoded shRNA and performed RNA-seq analysis. The RNA-seq results revealed at least two notable gene clusters (Cluster 4 and 5; Figure 1A) in terms of their dependence on HIRA and PML. Cluster 4 was comprised of genes whose expression decreased in senescent cells and were unaffected by HIRA and PML knock down (Figure 1A,B). Based on gene ontology analysis and pathways analysis, the top predicted upstream regulators of cluster 4 (Figure 1C), including E2F family members E2F4 and E2F1, MYC, RB1, CCND1, TP53, CDKN1A and CDKN2A, were associated with cell cycle pathways (Figure 1C, 1D). Cluster 4 primarily consisted of genes that were related to promoting proliferation^31^ (Figure 1E). This suggests that HIRA and PML are not necessary for the proliferation arrest observed in senescent cells. These findings were further confirmed by SA-β-Gal and EdU assays (Figure 1F,G).

**Figure 1:**
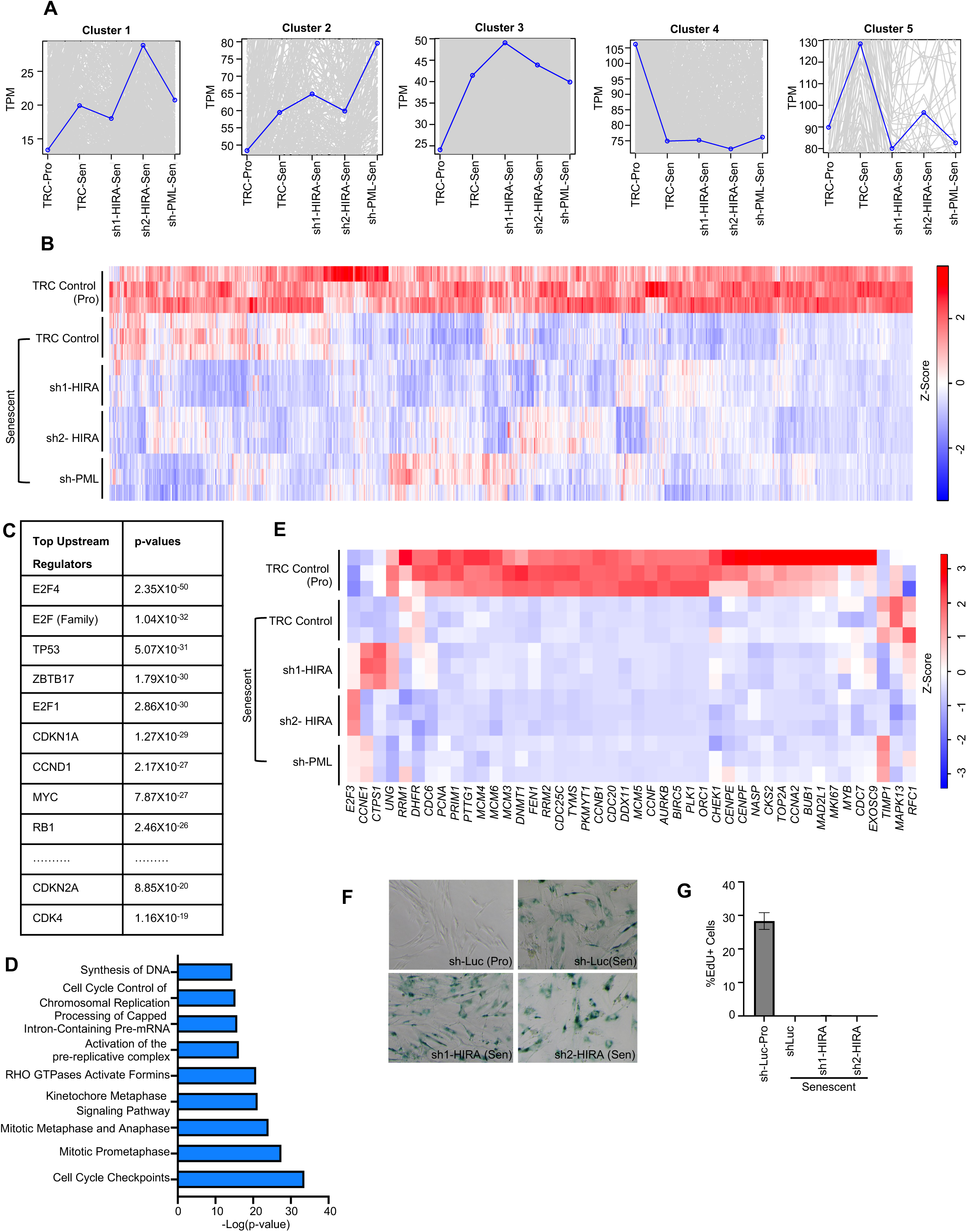
HIRA and PML are not essential for senescence-associated proliferation arrest. Senescence was induced in IMR-90 cells using 50 µM Etoposide for 24 hours. Cells were harvested for RNA-seq analysis 7 days post-treatment. (A) Clustering analysis of RNA-seq data reveals distinct gene expression patterns in five different clusters. (B) Heatmap of RNA-seq analysis showing the expression of Cluster 4 genes. (C) Gene ontology analysis by IPA identifies top upstream regulators in Cluster 4. (D) IPA canonical pathway analysis of Cluster 4 genes showed significant enrichment for Cell cycle pathways. (E) Heatmap representation of proliferation genes expression^31^. (F) Representative image of SA-β-Gal assay of control (shLuc) and HIRA-deficient (sh1,2-HIRA) senescent cells. (G) EdU assay was performed by treating cells with 10 μM EdU for 4 hours. In (B, E), heatmap color intensity represents the z-score calculated for each gene using TPM values, with red indicating high expression and blue indicating low expression. TRC Control and sh-Luc are the control shRNAs. Data shown in (A-E) represents three biological replicates. (G) represents three technical replicates.

Cluster 5 was comprised of genes that increased in senescence in a manner that depended on HIRA and PML (Figure 1A, 2A). Gene ontology analysis revealed that the top predicted upstream regulators of Cluster 5, including TNF, IL1A, IL1B, IL17A, LPS, NF-κB complex, RELA (NF-κB subunit p65) and IFNγ, were indicative of inflammatory pathways including interleukins, HMGB1, cGAS-STING signaling pathways (Figure 2B,C). These findings strongly suggest that HIRA and PML are necessary for the expression of the SASP, which we confirmed using NanoString nCounter technology, real-time qPCR, and western blotting of IL8 (Figure 2D-H). Furthermore, we observed that HIRA is essential for maximal SASP expression in various models of senescence, such as IR-induced senescence and oncogene-induced senescence (Figure S4A,B).

**Figure 2:**
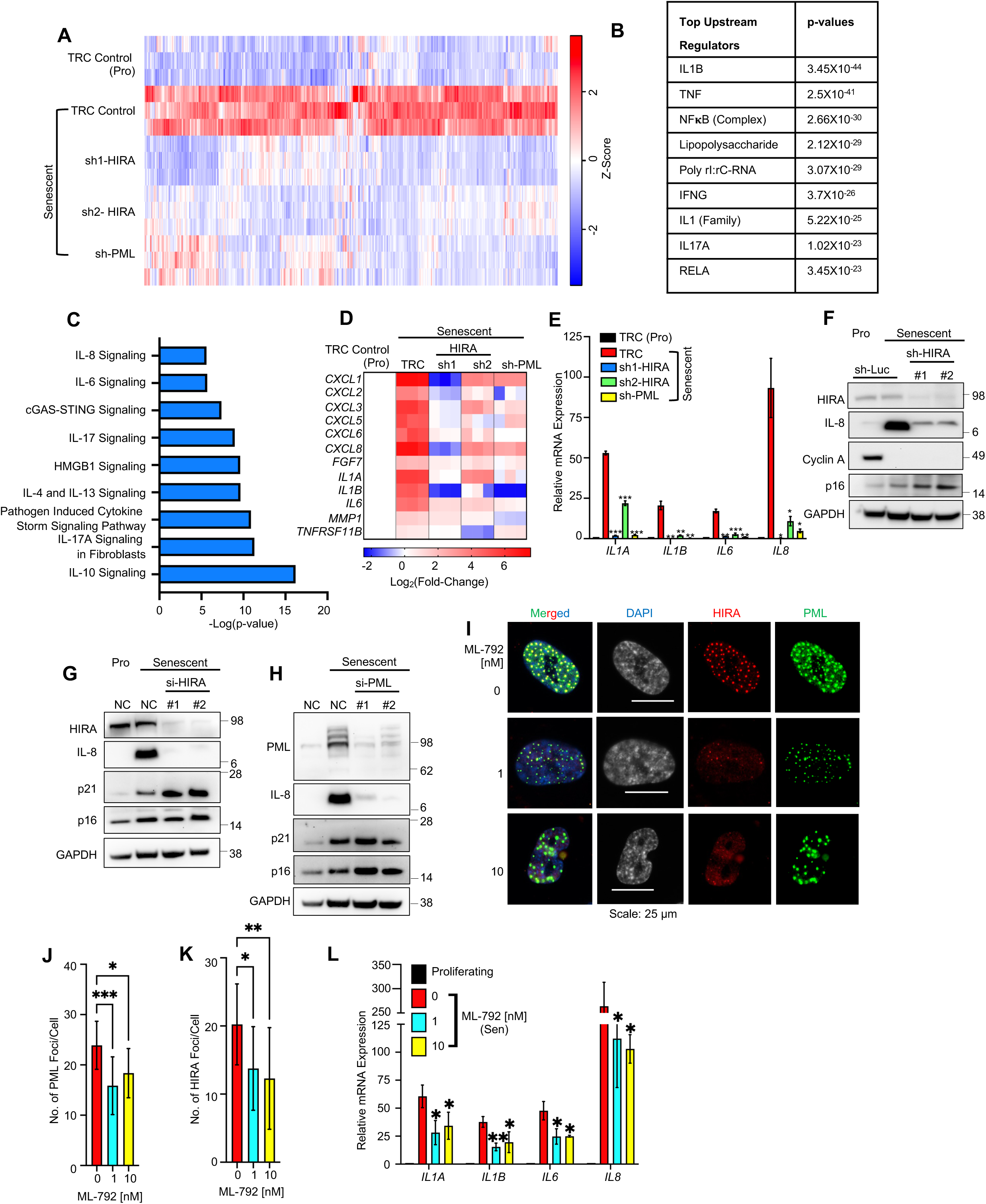
HIRA and PML are required for SASP expression. (A) Heatmap of RNA-seq analysis showing the expression of Cluster 5 genes (Figure 1A). (B) Top upstream regulators of Cluster 5 identified by IPA analysis. (C) IPA canonical pathway analysis of Cluster 5 genes showed significant enrichment for inflammatory pathways. (D) Heatmap of NanoString analysis and (E) Real-time qPCR analysis showing the expression of SASP genes in control (TRC) and HIRA (sh1,sh2), and PML (sh-PML) deficient cells. (F) Western blot analysis of IL-8 as a SASP marker, and Cyclin A as proliferation marker and p16 as senescence markers, was performed in HIRA-deficient senescent cells under the similar conditions as the RNA-seq experiment. sh-Luc is the control shRNA targeting luciferase. (G,H) Further validation of RNA-seq results through Western blot analysis following the knockdown of HIRA and PML using siRNAs. IL-8 was used as a marker for the SASP, while p21 and p16 were used as markers for senescence. Senescence was induced with 25 µM Etoposide for 24 hours, and siRNAs were transfected the following day. Cells were harvested for RNA-seq analysis 7 days post-Etoposide treatment. “NC” denotes the negative control siRNA. (I) Representative immunofluorescence image showing the localization of HIRA and PML in senescent cells in the presence of the drug ML-792. (J,K) Quantification of the number of PML and HIRA foci in senescent cells in the presence of drug ML-792, respectively. (L) Real-time qPCR analysis of SASP genes in the presence of ML-792 (0,1,10 nM). For (I-L) senescence was induced using 50 µM Etoposide for 24 hours. ML-297 was added the next day, and the media, replenished with the drug, was changed every 2 days. Cells were harvested and fixed for qPCR and immunofluorescence analysis, respectively, 7 days post-Etoposide treatment. In (A), heatmap color intensity represents the z-score calculated for each gene using TPM values, with red indicating high expression and blue indicating low expression. Data shown in (A-D), and (L) represents three biological replicates. Figures (E,J,K,L) represent the mean ± SD. The p-values were calculated using an unpaired two-tailed Student’s t-test (E,L), and using one-way ANOVA with Dunnett’s multiple comparisons test (J,K). The images were captured automatically using Nikon motorized platform. The values were calculated using Nikon NIS-Element software from 3 different wells with multiple fields. (∗∗∗) p < 0.001; (∗∗) p< 0.01; (∗)p< 0.05.

To investigate a role for HIRA/PML organization in the form of PML NBs for SASP expression, we conducted an experiment using a low dose of the SUMO Activating Enzyme (SAE) inhibitor ML-792 to disrupt the organization of PML NBs, as previously reported^32^. ML-792 impaired PML focal organization (Figure 2I,J), and dispersed the localization of HIRA (Figure 2I,K). Additionally, we made an interesting observation in senescent cells, whereby a low abundance of a higher apparent molecular weight form of HIRA was present in senescent cells (Figure S4C). Inhibition of SUMOylation using ML-792 led to the disappearance of this isoform (Figure S4C). Knocking down SUMO-conjugating enzyme UBC9 with UBE2I siRNA also resulted in the disappearance of this polypeptide (Figure S4D). This isoform was also absent in senescent cells lacking PML (Figure S4E). These findings suggest that HIRA undergoes SUMOylation within PML NBs in senescent cells. Our results also demonstrated that ML-792 was capable of suppressing the SASP (Figure 2L), supporting the notion that the proper organization of HIRA/PML is necessary for maximal SASP expression.

In summary, our results demonstrate that while HIRA and PML are not essential for the arrest of proliferation, they play a crucial role in SASP expression. Moreover, the use of a drug-like inhibitor that disrupts the functional integrity of PML NBs by blocking SUMOylation^32^ also suppressed the SASP, suggesting that SASP may depend on the co-localization of HIRA and PML in PML NBs.

### Localization of HIRA in PML NBs is tightly linked to SASP expression

Based on these findings, we aimed to more directly investigate whether the co-localization of HIRA and PML is required for the expression of SASP. To address this, we utilized a mutant form of HIRA called ⊿HIRA (520-1017) (referred to as ⊿HIRA), which we previously showed to localize to PML NBs, disrupt the localization of endogenous HIRA to PML NBs, and exhibit dominant-negative inhibitory activity over HIRA function^22^. We successfully expressed ⊿HIRA and confirmed its localization to PML NBs (Figure 3A,B). Furthermore, using an antibody that detects endogenous full length HIRA but not truncated ⊿HIRA^22^, we observed that ⊿HIRA suppressed the localization of endogenous HIRA to PML NBs (Figure 3C-F), while the number of PML NBs remained unchanged (Figure 3E). Importantly, ⊿HIRA demonstrated a significant blockade of the expression of various inflammatory cytokines, including IL1A, IL1B, IL6, and IL8, as determined by quantitative PCR (qPCR) and western blot (WB) analysis (Figure 3G,H), without affecting downregulation of cyclin A and upregulation of p16 and p21 (Figure 3H). To further validate these findings, we performed RNA-seq analysis on senescent cells expressing ⊿HIRA and control cells. The results revealed a comparable number of differentially expressed genes (DEGs) in senescent versus proliferating cells in both the ⊿HIRA and wild-type samples, with 3821 genes upregulated and 3340 genes downregulated in the ⊿HIRA cells, and 3549 genes upregulated, and 3180 genes downregulated in the wild-type cells (Figure S5A, B).

**Figure 3:**
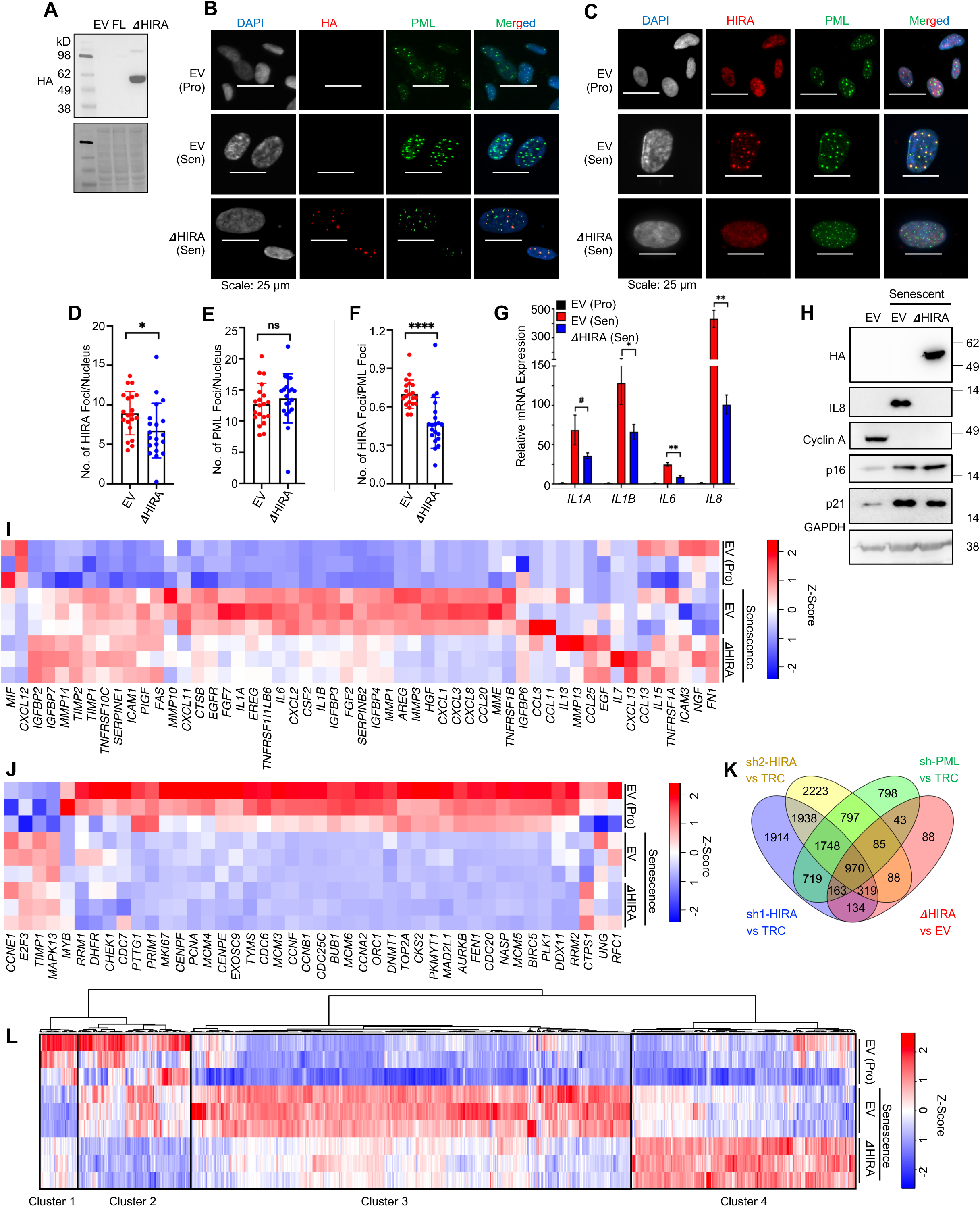
Localization of HIRA to PML NBs is tightly linked to SASP expression. (A) Expression of empty vector (EV), wild-type HIRA (FL), and ΔHIRA(520-1017) in senescent IMR-90 cells by western blot. The predicted molecular weight of HIRA (FL) is 112 kDa, while ΔHIRA is expected to be 55 kDa. (B) Representative immuno-fluorescence images showing ΔHIRA expression in senescent cell nuclei, stained with HA antibody. (C) Representative immuno-fluorescence images illustrating the localization of endogenous HIRA in control (EV) and ΔHIRA cell lines. The WC119 antibody used to detect endogenous HIRA does not recognize ΔHIRA (520-1017)^22^. (D-F) the number of HIRA foci, PML foci, and their ratio, respectively. Data represents the mean ± SD. The values were automatically calculated using Nikon-NIS software from three different wells with multiple fields in an unbiased manner. (G) Real-time qPCR analysis showing the expression of SASP genes in control (EV) and ΔHIRA senescent cells. Data shown represents the mean ± SD of n = 3 biological replicates. (H) Immunoblot analysis of IL8 as a SASP gene, Cyclin A as a proliferation marker, and p21, p16 as senescent markers in control (EV) and ΔHIRA senescent cells. (I,J) Heatmap of RNA-seq analysis displaying the expression of SASP gene^2^ (I); and proliferation genes^31^ (J) in proliferating control (EV (Pro)), senescent control (EV), and senescent ΔHIRA cells, 8 days after Etoposide treatment. Senescence was induced in IMR-90 cells using 50 µM Etoposide for 24 hours. (K) Venn diagram depicting differentially expressed genes (DEGs) in ΔHIRA, two HIRA-knockdown, and one PML-knockdown senescent cells, generated using VENNY2.1. (L) Heatmap of the 864 genes in proliferating control (EV (Pro)), senescent control (EV), and senescent ΔHIRA cells, which are altered in the same direction by both HIRA shRNAs and PML shRNA. In I, J, L, color intensity represents the z-score calculated for each gene using TPM values, with red indicating high expression and blue indicating low expression. Data shown represents n = 3 biological replicates. For data (D-G), the p-values were calculated using an unpaired two-tailed Student’s t-test. (∗∗∗) p < 0.001; (∗∗) p < 0.01; (∗) p < 0.05, (#) p<0.1.

Intriguingly, 1145 and 745 genes were down and upregulated respectively by ⊿HIRA in senescent cells (Figure S5A,B). Similar to HIRA knock down, ⊿HIRA inhibited SASP^2^ genes but had no impact on proliferation-related genes^31^ (Figure 3I, J). Remarkably, 35.7% of the genes (970/ (970+1748) X 100) affected by both HIRA shRNAs and PML shRNA were also altered by ΔHIRA (Figure 3K). Among the 970 DEGs, the expression of 864 genes (89%) was altered in the same direction by both HIRA shRNAs, PML shRNA, and ΔHIRA. The fold enrichment of these 864 DEGs over random overlap was calculated as 25.9, with a p-value of less than 2.2X10^-308^ (two-tailed Fisher exact test). These 864 subdivided into four distinct clusters (Figure 3L, S5C). The largest cluster, Cluster 3 (nearly 50% of the 864 genes), was comprised of genes whose expression increases in senescence and is decreased on expression of ⊿HIRA. These genes were primarily comprised of inflammatory pathways, including cGAS-STING signaling (Figure S5C). Taken together, these results obtained with ⊿HIRA, in conjunction with those from the SUMOlyation inhibitor, strongly suggest that the proper spatial configuration of HIRA and PML NBs in senescent cells is crucial for the expression of SASP.

To further verify the involvement of HIRA localization within PML NBs in SASP expression, we employed two additional methodologies. Firstly, after SP100 depletion, we observed that HIRA did not aggregate into nuclear foci during interferon stimulation, as reported earlier^33^ (data not shown). Similarly, knockdown of SP100 led to a notable reduction in HIRA foci in senescent cells, without affecting PML foci (Figure 4A-F). Notably, the mRNA level of HIRA expression remained constant (data not shown). Importantly, we observed a decrease in SASP expression upon SP100 depletion (Figure 4G-I). As a second approach, we took advantage of the fact that we previously established that glycogen synthase kinase 3β (GSK-3β) phosphorylates HIRA on S697, facilitating its accumulation in PML NBs during senescence initiation^34^. Here, we employed LY2090314, a GSK-3β inhibitor currently undergoing clinical trials, to block GSK-3β activity. Remarkably, while the number of PML or SP100 foci remained unchanged (Figure 4J-M), the number of HIRA foci within PML NBs markedly decreased (Figure 4J,K,N), correlating with a reduction in SASP expression (Figure 4O), in line with the other approaches for disrupting HIRA/PML colocalization. Based on the data presented from multiple approaches - knock down of HIRA, PML and SP100, ectopic expression of ⊿HIRA, and inhibiting SUMOylation and GSK-3β, we can conclude that the localization of HIRA in PML NBs is tightly associated with and likely causally important for SASP expression.

**Figure 4:**
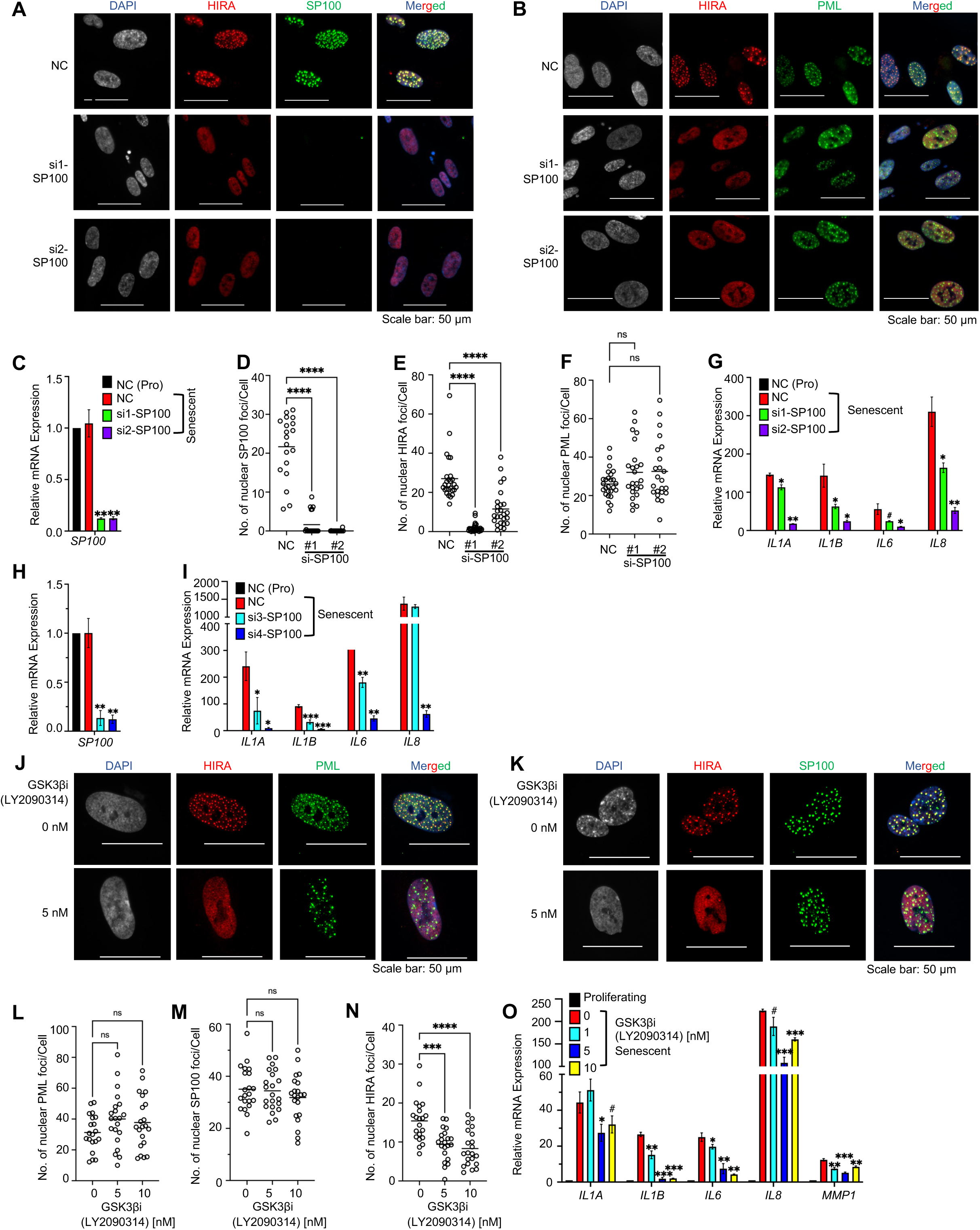
HIRA localization to PML NBs and SASP expression are SP100 and GSK3**β** dependent. (A) Representative immunofluorescence image showing the dispersion of HIRA foci in senescent cells in the absence of SP100. (B) Representative immunofluorescence image showing the localization of HIRA and PML in senescent cells in the absence of SP100. (C) Relative *SP100* mRNA expression and (D) the number of SP100 foci based to calculate the efficiency of *SP100* siRNA (#1,#2). (E,F) Quantification of the number of HIRA and PML foci in senescent cells in the absence of SP100, respectively. (G) Real-time qPCR analysis of SASP genes in the absence of SP100 using si-RNA (#1,#2). (H,I) Relative *SP100* and SASP expression in the absence of SP100 using si-RNA (#3,#4). Senescence was induced using 25 µM Etoposide for 24 hours. The next day, 20 nM siRNAs were transfected. Cells were harvested and fixed for qPCR and immunofluorescence analysis 7 days post-Etoposide treatment. (J,K) Representative immunofluorescence image showing the localization of HIRA, PML and SP100 in senescent cells in the presence of the drug LY2090314, a GSK-3β inhibitor. (L,M,N) Quantification of the number of PML, SP100, and HIRA foci in senescent cells in the presence of the drug LY2090314, respectively. (O) Real-time qPCR analysis of SASP genes in the presence of the drug LY2090314 (0,1,5,10 nM). Senescence was induced using 50 µM Etoposide for 24 hours. LY2090314 was added the next day, and the media, replenished with the drug, was changed every 2 days. Cells were harvested and fixed for qPCR and immunofluorescence analysis, respectively, 7 days post-Etoposide treatment. For (D-F,L-N), the images were captured automatically using Nikon motorized platform. The values were calculated using Nikon NIS-Element software from 3 different wells with multiple fields. The p-values were calculated using an unpaired two-tailed Student’s t-test (C,G-I,O), and using one-way ANOVA with Dunnett’s multiple comparisons test (D-F,L-N). (∗∗∗) p < 0.001; (∗∗) p< 0.01; (∗)p< 0.05.

### HIRA regulates SASP through activation of NF-***K***B pathway independent of its H3.3 deposition role

We set out to define the role of HIRA and PML in expression of SASP. Since NF-κB serves as the master regulator of SASP expression, we initially examined whether HIRA is necessary for NF-κB activation. We observed a decrease in NF-κB luciferase reporter activity in senescent cells depleted of HIRA (Figure 5A). This led us to propose that HIRA/PML might be essential for expression of SASP genes through a requirement for HIRA for transcription-coupled deposition of histone H3.3 at SASP. To test this hypothesis, we asked whether knockdown of HIRA would also impede the expression of SASP genes induced by stimuli other than senescence. We treated proliferating and senescent cells, with or without HIRA depletion through shRNA knockdown, with recombinant IL1A (rIL1A). To our surprise, and contrary to our hypothesis, we found that rIL1A effectively induced IL8 expression in both proliferating and senescent cells, irrespective of HIRA knockdown. In other words, the addition of rIL1A restored SASP expression in senescent cells lacking HIRA (Figure 5B,C), suggesting that HIRA-dependent transcription-coupled deposition of histone H3.3 is not essential for expression of these genes. To further investigate the requirement for deposition of histone H3.3 at SASP genes in senescent cells, we used siRNA to deplete H3.3 and assessed the expression of SASP genes such as *IL8*, *IL1A*, *IL1B*, and *IL6*. Remarkably, knockdown of H3.3 resulted in increased expression of these SASP genes compared to control senescent cells (Figure 5D,E). Consequently, we concluded that the requirement for HIRA/PML in SASP expression does not involve HIRA’s role in transcription-coupled deposition of histone H3.3. We reasoned that HIRA and PML must be necessary for SASP expression upstream of gene transcription. Therefore, we examined the impact of HIRA knockdown on p65(RelA) nuclear translocation, phosphorylation, and other markers of NF-kB activation. Our results revealed that HIRA depletion in senescent cells led to suppressed p65(RelA) nuclear translocation (Figure 5F). Additionally, HIRA was crucial for the activation of the entire canonical NF-κB pathway, as evidenced by p65(RelA) phosphorylation, IKBα phosphorylation and degradation, and IKKβ/α phosphorylation (Figure 5G). Furthermore, PML was also essential for p65(RelA) activation (Figure 5H). In conclusion, inactivation of HIRA does not directly block expression of NFkB target genes; instead, a non-canonical/H3.3 independent HIRA function is required for signaling to activate NF-κB in senescent cells and induce SASP expression.

**Figure 5:**
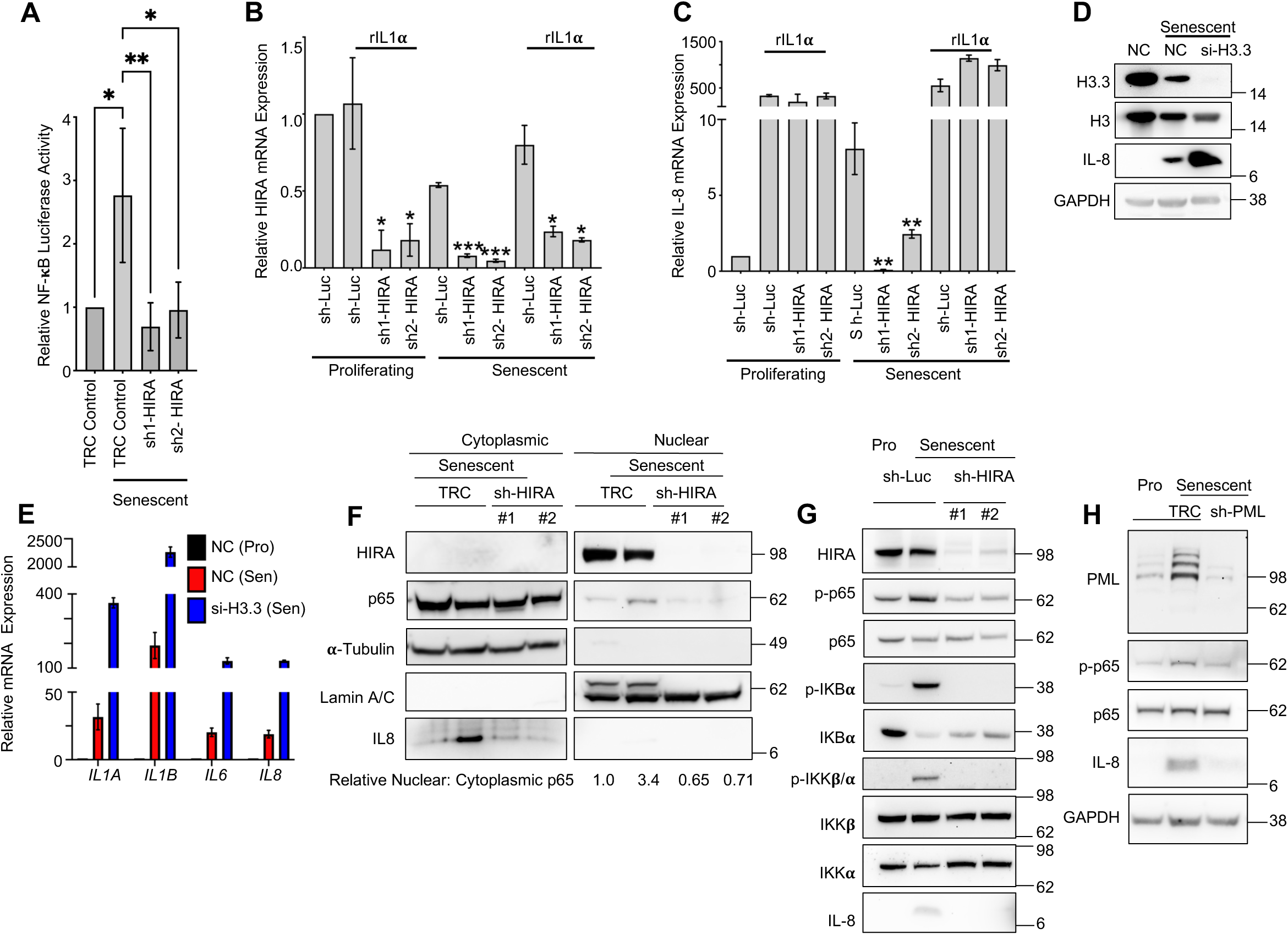
HIRA regulates SASP through activation of the NF-**K**B pathway independent of its H3.3 deposition function. (A) The relative luminescence was measured from the lysate obtained from lenti-NF-κB-luc/GFP reporter IMR-90 cells. The luciferase activity was normalized to the mean fluorescence of GFP. (B,C) Real-time qPCR analysis was conducted to measure the expression of HIRA as an indicator of knockdown efficiency and IL-8 as SASP gene in proliferating and senescent control cells (sh-Luc control) and HIRA-deficient cells (sh1, sh2-HIRA) after treatment with recombinant IL1α (r IL1α 20 ng) for 24 hours. Senescence was induced using 50 µM Etoposide for 24 hours. On day 7 post-Etoposide treatment, rIL1α was added without changing the medium, and cells were harvested 24 hours after the addition of rIL1α. (D) Immunoblot analysis was performed on proliferating control cells (NC), senescent control cells (NC), and H3.3 knock-down cells. (E) Real-time qPCR analysis was conducted to assess the expression of SASP genes in control (NC) and H3.3-deficient senescent cells (si-H3.3). For (D, E), senescence was induced using 25 µM Etoposide for 24 hours. The next day, 20 nM siRNAs were transfected, and the cells were harvested on day 7 post-Etoposide treatment. (F) The nuclear translocation of p65 was analyzed by cellular fractionation followed by western blotting. α-Tubulin was used as a cytoplasmic marker, and Lamin A/C was used as a nuclear marker. The ratio of nuclear p65 to cytoplasmic p65 is given, normalized to the proliferating control. (G) The canonical NF-κB pathway was analyzed by immunoblotting in HIRA-depleted cells. (H) Immunoblot analysis was performed to examine p65 phosphorylation in PML-depleted cells. For (F-H), senescence was induced using 50 µM Etoposide for 24 hours and the cells were harvested on day 7 post-Etoposide treatment. Both TRC and sh-Luc were used as control samples. Data shown in (A-C), and (E) represents the mean ± SD of n = 3 biological replicates. The p-values were calculated using an unpaired two-tailed Student’s t-test. (∗∗∗) p < 0.001; (∗∗) p< 0.01; (∗) p< 0.05.

### HIRA and PML are required for cGAS-STING-TBK1 signaling

Numerous upstream factors have been implicated in the activation of NF-κB and SASP in senescent cells, including mTOR, GATA4, CEBPB, DDR, and cGAS/STING. However, upon closer analysis, we failed to observe a consistent effect of HIRA/PML on regulation of certain effectors involved in SASP, such as mTOR (reflected in p70S6K phosphorylation), GATA4, p38MAPK, CEBPB, and γH2AX^8^ (Figure S6A,B). We hypothesized that HIRA is essential for the activation of the cytoplasmic CCF-cGAS-STING pathway, which in turn promotes NF-κB activation and SASP expression. Our observations revealed that both HIRA and PML play essential roles in TBK1 activation (Figure 6A, B), STING dimerization (Figure 6C), and cGAMP production (Figure 6D). Consistent with a role upstream of cGAS/STING activation, HIRA, primarily known for its nuclear localization, exhibited colocalization with cGAS in the cytoplasmic foci (CCF) (Figure 6E-G). Although HIRA was not necessary for CCF formation (data not shown), the absence of HIRA resulted in compromised enrichment of cGAS at the CCF (Figure 6H, I). Notably, we did not observe any consistent changes in the expression of cGAS in cells lacking HIRA (data not shown). These findings suggest that HIRA regulates the cGAS-STING-TBK1-NF-κB axis through an unknown mechanism linked to regulated enrichment of cGAS at CCF.

**Figure 6:**
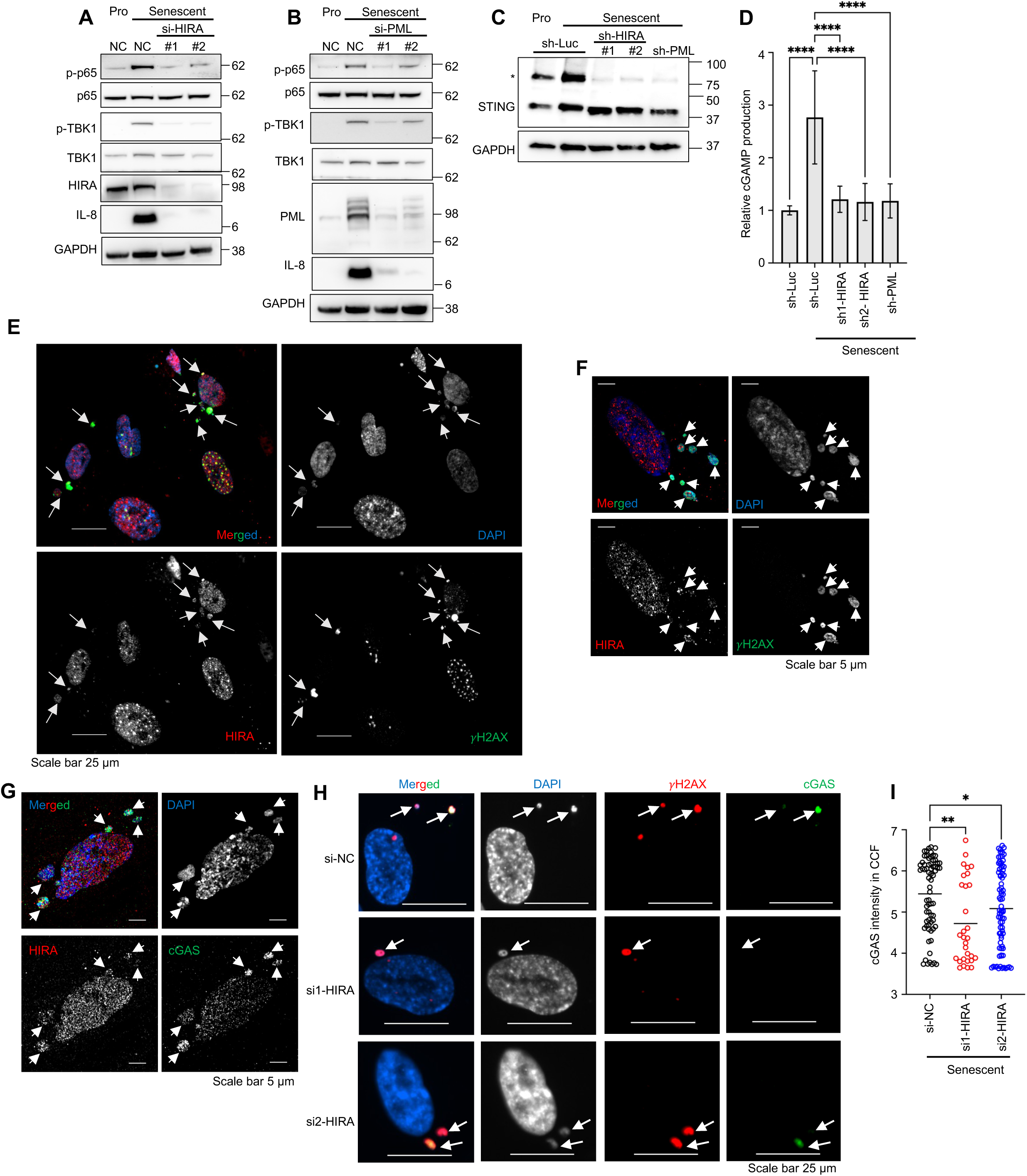
HIRA and PML are required for cGAS-STING-TBK1 signaling. (A-C) Cell lysates were subjected to immunoblotting. STING blot in (C) was performed under non-reducing condition. * indicates STING dimer. (D) cGAMP synthesis was assessed using the ELISA method after 8 days of etoposide treatment. Data represent the mean ± SD of n = 6 (3 biological replicates from two independent experiments). The p-values were calculated using one-way ANOVA with Dunnett’s multiple comparisons test. (E) Epifluorescence image of localization of HIRA in CCF. (F,G) Representative images of Airyscan Super-Resolution microscopy of HIRA in CCF. (H,I) Cells were stained for DAPI, γH2AX, and cGAS in control (NC) and HIRA-deficient cells (si1,2-HIRA). The cGAS intensity in individual CCF (DAPI and γH2AX positive foci in cytoplasm) was calculated. Each data point represents an individual CCF. The values were calculated using Nikon NIS-Elements software from 3 different wells with multiple fields and were log-transformed. For (A, B, H, I), senescence was induced using 25 µM Etoposide, and for (C-G), 50 µM for 24 hours. The cells were harvested on day 7 post-treatment. The p-values were calculated using an unpaired two-tailed Student’s t-test. (∗∗∗∗) p < 0.001;(∗∗∗) p < 0.001; (∗∗) p< 0.01; (∗) p< 0.05.

### HIRA physically interacts with autophagy regulator p62, a repressor of SASP

To investigate the non-canonical cytoplasmic role of HIRA in the expression of SASP, we conducted a study to identify potential effectors responsible for this function. We employed immunoprecipitation followed by mass spectrometry (IP-MS) on both proliferating and senescent cells to discover novel candidate effectors associated with SASP (Supplementary Excel File 1). Subsequently, we validated several candidates through immunoprecipitation followed by Western blotting (IP-WB), namely FHL2, LUZP1, PML, NBR1, SQSTM1 (p62), SP100, and MAGED1 (Figure 7A-C). Previously, we demonstrated the striking partial localization of p62 in CCF^10^. In this study, we further confirmed the interaction between HIRA and p62 through p62 immunoprecipitation (Figure 7D). Moreover, we observed the co-localization of p62 with HIRA in PML NBs in senescent cells (Figure 7E). Through siRNA knockdown experiments, we discovered that p62 functions as a robust negative regulator of SASP expression (Figure 7F). To validate this finding, we ectopically expressed p62, resulting in the potent suppression of SASP expression (Figure 7G). Ectopic expression of p62 not only dampened TBK1 and p65 activation and IL8 expression (Figure 7H), but also reduced cGAS levels in senescent cells (Figure 7I) and suppressed STING dimerization (Figure 7J), without affecting the expression level of HIRA (Figure S7A). Conversely, although knock down of p62 similarly did not affect HIRA levels (Figure S7B), we observed an increase in cGAS levels in p62-depleted senescent cells compared to control senescent cells (Figure S7C,D). These results implicate p62 as a negative regulator of SASP, acting on the cGAS/STING-TBK1-NFkB pathway, potentially antagonistic to HIRA. Indeed, previous studies have demonstrated a role for p62 in autophagic degradation of cGAS and STING ^35,36^. Confirming an antagonistic relationship between HIRA and p62 in regulation of SASP, knockdown of p62 rescued SASP expression in HIRA-deficient cells, and knockdown of HIRA decreased SASP in p62-deficient cells (Figure 7K). In sum, HIRA physically interacts with p62, a negative regulator of SASP. HIRA and p62 antagonistically regulate the SASP Similar to HIRA, p62 localized to CCF (Figure S8A)^10^, and HIRA and p62 physically interacted in senescent cells (Figure 7D). We investigated whether PML is necessary for this interaction. However, in the absence of PML, we observed that the physical interaction between HIRA and p62 remained unchanged, as demonstrated by immunoprecipitation followed by Western blotting in PML-depleted cells (Figure S8B,C). However, the absence of PML caused mislocalization of both HIRA and p62 and reduced the number of dual HIRA and p62 foci in the nucleus (Figure 7L-N and S8D-F). Consequently, the absence of PML altered the spatial relationship between p62 and HIRA (Figure 7O).

**Figure 7:**
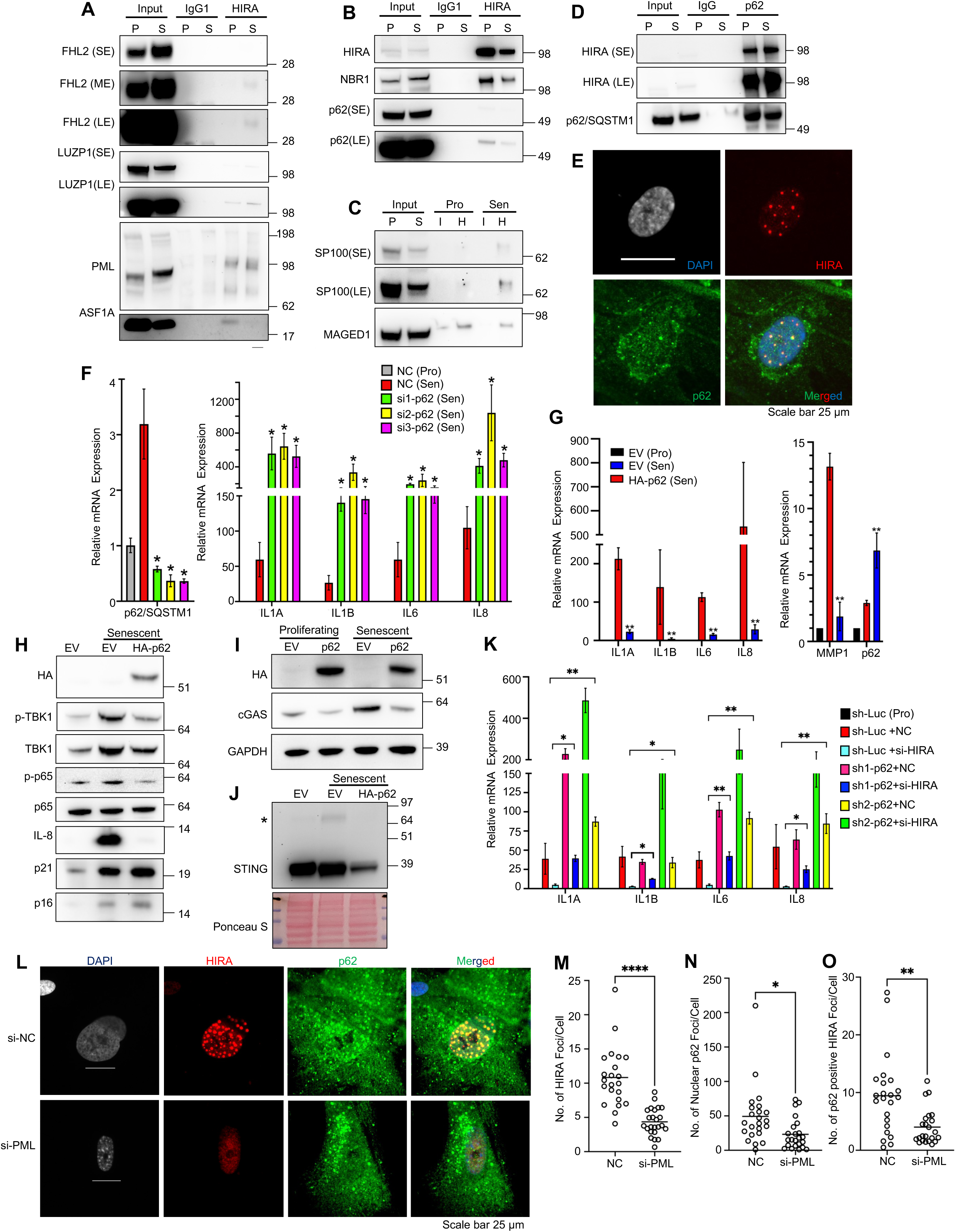
HIRA physically interacts with autophagy regulator p62 to regulate SASP expression. (A-C) Validation of HIRA interactor proteins by western blotting, obtained from IP-MS. P, proliferating; S, senescent; I, IgG1; H, antibody against HIRA (D) Immunoprecipitation of p62 interactor proteins followed by immunoblotting for HIRA. Short exposure (SE) and long exposure (LE) were performed. (E) Representative image showing the localization of p62 with HIRA in PML NBs of senescent cells. (F) Real-time qPCR analysis showing the expression of SASP genes in control (NC) and p62-deficient cells (si1, si2, and si3). (G-I) Real-time qPCR analysis showing the expression of SASP genes (G) and immunoblot analysis of cytoplasmic fraction of senescent empty vector (EV) and HA-p62 expressing cells (H, I). (J) STING immunoblot was performed under non-reducing conditions. * indicates STING dimer. (K) Real-time qPCR analysis showing the expression of SASP genes. IMR-90 cells were first infected with lentivirus expressing shRNA against p62 (sh1 and sh2) and control (sh-Luc), and puromycin (1μg/mL, 2 days) selected, as indicated. The day following etoposide (25 μM) treatment, the cells were transfected with negative control siRNA (NC) or si-HIRA, as indicated. (L) Representative image showing the localization of p62 with HIRA in the presence and absence of PML NBs in senescent cells. (M-O) The random images were captured automatically using Nikon motorized platform. The values were calculated using Nikon NIS-Element software from 3 different wells with multiple fields. Data in (F) and (K) represent the mean ± SD of n = 3 biological replicates. (G) represents three biological replicates from two independent experiments (n=3X2). For (A-E,H-J), senescence was induced using 50 µM Etoposide, and for (F,K-O) 25 µM for 24 hours. The cells were harvested on day 7 post-treatment. For data in (F), (G), and (K), the p-values were calculated using an unpaired two-tailed Student’s t-test. (∗∗∗∗) p < 0.0001; (∗∗∗) p < 0.001; (∗∗) p < 0.01; (∗) p < 0.05.

## Discussion

In this study, we have demonstrated that the histone chaperone HIRA physically interacts with the PML protein and SP100, and localizes to PML NBs in senescent cells. HIRA is also required for SASP expression, without affecting cell proliferation arrest. By multiple approaches, we showed that HIRA localization to PML NBs is tightly linked to SASP. Although off target effects might account for the effects on SASP, this seems unlikely given the diversity of approaches employed. Furthermore, our investigation revealed that HIRA exerts a non-canonical non-histone chaperone role in the regulation of SASP, independently of its role in H3.3 deposition. In this process, HIRA directly interacts with the autophagy cargo receptor p62, which functions as a negative regulator of SASP. The p62 protein is able to downregulate cGAS. Since both cGAS and p62 are enriched in CCFs, it is probable that p62 mediates the degradation of cGAS within the CCF^35^, thereby reducing the interaction between CCF and cGAS. Additionally, it can be speculated that the physical interaction between HIRA and p62 somehow restores cGAS within the CCF. This subsequent activation of the cGAS-STING-TBK1-NF-κB signaling axis leads to the expression of SASP. In the absence of PML, HIRA and p62 foci become diffused within the nucleus, potentially disrupting the normal spatial and regulatory relationship between HIRA and p62.

Multiple reports describe the induction of PML during senescence, suggesting its correlation with the inhibition of E2F gene expression, which is proposed to induce senescence^27-29^. However, in our observations, although PML is induced at the protein level in senescent cells, the number and intensity of PML foci increase, but it is not essential for proliferation arrest. Moreover, in the absence of PML, E2F target gene expressions remain dampened. This suggests that although PML induction and E2F target gene repression are correlated, PML is not essential in all contexts for repression of E2F target genes.

It is well established that PML NBs are associated with the interferon system and play a crucial role in anti-viral defense^37^. The cGAS/STING pathway is also involved in anti-viral defense. Upon binding to viral double-stranded DNA, cGAS initiates a tightly regulated signaling cascade involving the adapter STING^13,38^. This cascade triggers various inflammatory effector responses by activating transcription factors including IRF3 and NF-κB^13,38^. Previous research conducted by our laboratory has demonstrated that HIRA is also engaged in the detection of foreign viral DNA^39^. Moreover, HIRA actively participates in antiviral functions by promoting the expression of cellular genes associated with innate immunity and a diverse array of interferon-stimulated genes (ISGs)^33^. Here, we have shown that PML is also involved in cGAS/STING signaling and expression of SASP in response to another type of cytoplasmic DNA, CCF. Here we have also observed the presence of HIRA within CCFs. In the process of senescence, as well as during DNA virus infection, DNA transfection, and interferon treatment, HIRA accumulates in PML NBs^21,39^. Thus, HIRA responds to foreign and cytoplasmic DNAs in a way that links it to PML. PML and HIRA’s localization to PML NBs are required for expression of SASP. Therefore, the shared involvement of HIRA, PML, cGAS, and STING in cellular senescence and intrinsic antiviral immunity suggests that “seno-viral” signaling acts as a nexus linking cellular innate immunity and cell senescence.

PML within nuclear bodies (NBs) has been suggested to finely regulate various processes by facilitating posttranslational modifications, particularly SUMOylation, of partner client proteins, including PML itself^23,40^. SUMOylation of PML is a prerequisite for NB formation^41^ and SUMOylation of partners can lead to partner sequestration, activation, or degradation, thereby influencing a wide range of cellular functions including senescence^23,40,42,43^. The overexpression of SUMO has been found to induce senescence, as increased SUMOylation of specific target proteins can lead to premature cellular senescence^42,43^. Our preliminary data suggests that in addition to PML, HIRA is also likely to undergo SUMOylation. SUMOylation requires the presence of a consensus SUMOylation motif within the target protein. By performing bioinformatic analysis, we have identified several potential SUMOylation sites within the HIRA protein, such as K92, K617, K809, and K843. However, further investigation is needed to confirm the occurrence of HIRA SUMOylation and determine the sites and function of SUMOylation during senescence. Based on our findings, we propose that HIRA translocates to PML NBs in senescent cells, perhaps for the purpose of SUMOylation, which significantly influences its functionality. It is noteworthy that the absence of HIRA does not alter the number of PML NBs, yet it leads to a decrease in SASP expression. Hence, in the HIRA/PML axis, HIRA plays a pivotal role in expressing SASP, while PML NBs appear to act as a molecular platform for spatial interaction of HIRA with other proteins, such as p62, and post-translational modification, SUMOylation in particular. Considering the increasing recognition of the significance of senescence, SUMOs, and PML in cancer development and therapeutic response^43^, this finding represents an exciting development, warranting thorough exploration and study.

Recent research has emphasized the importance of p62 in senescence and SASP regulation. The p62 protein acts as a negative regulator of SASP in GATA4-mediated NF-κB activation^44^, which aligns with the findings of our study. Furthermore, p62 functions as a cargo receptor for the degradation of proteins in the inflammation pathway. During viral infection or dsDNA stimulation, p62-mediated autophagy facilitates the degradation of cGAS, STING, and IRF3, which are key components of the cGAS-STING pathway^35,36,45,46^. Consequently, p62 exerts negative control over the cGAS-STING pathway, thereby regulating type I IFN signaling during virus infection^35,36,46^. However, in the case of dsDNA stimulation, STING is specifically degraded through a p62-dependent mechanism^36^. Although p62 does not directly bind to cGAS-associated dsDNA, it predominantly colocalizes with STING upon dsDNA transfection^36^. In contrast, p62 is abundant in cGAS-bound CCFs, suggesting p62 may mediate the degradation of cGAS associated with CCF, as suggested by p62 overexpression and depletion in senescent cells. While the exact role of p62 and cGAS degradation in this study is not fully elucidated, it is speculated that the physical and spatial interaction between p62, HIRA, and PML regulates the CCF-cGAS-STING-NF-κB axis in senescence. Ongoing and future studies will address these questions. The inclusion of p62 in both the HIRA/PML axis and cGAS-STING pathway in senescence and antiviral signaling provides additional support for the concept of “seno-viral” signaling and suggests an evolutionary connection between viral infection and senescence.

One limitation of this study is that HIRA has previously been reported to be required for SAHF formation in senescent cells. We have confirmed this (data not shown), aligning with our previous observations ^21,22^. A previous report showed that SAHF formation is required for SASP^47^, and we have confirmed this (data not shown), suggesting that the requirement for HIRA for SASP might reflect its requirement for SAHF. However, if a HIRA-SAHF-SASP pathway exists, it is poorly defined and its relationship to the HIRA-p62-cGAS-SASP pathway proposed here is unclear. Further investigations are required. A number of other limitations reflect unanswered mechanistic questions. Our immunofluorescence data showed colocalization of p62 and HIRA in CCF, and we have observed a physical interaction between these two proteins. This suggests they interact within CCF, yet isolating CCF to examine this interaction has not been possible for us. While p62 and HIRA antagonistically regulate SASP, potentially through cGAS in CCF, we have not dissected the exact mechanism involved. HIRA localization in PML NBs is tightly linked to SASP expression, likely influenced by posttranslational modifications such as phosphorylation and SUMOlyation of HIRA in PML NBs. However, we have not identified these modifications and confirmed their necessity. However, it is possible that HIRA regulates SASP through both PML-dependent and PML-independent mechanisms. Finally, SQSTM1 (p62) plays a crucial role in targeting ubiquitinated proteins for degradation through the autophagy pathway and previous studies have demonstrated a role for p62 in autophagic degradation of cGAS and STING ^35,36^. However, p62 is also implicated in the ubiquitin-proteasome system (UPS)^48^. Conceivably, p62 alternatively regulates SASP via control of cGAS/STING through the UPS. Ongoing and future studies will address these outstanding mechanistic questions.

In summary, this study points to a novel cytoplasmic role of HIRA in the cGAS-STING signaling pathway in senescent cells, and reveals a shared role for HIRA and PML in control of SASP, likely mediated by PML’s role in organizing the spatial configuration of HIRA and p62, antagonistic regulators of SASP. The findings strengthen the connection between cytoplasmic/foreign DNA sensing mechanisms in virus infection and senescence and highlights the potential therapeutic significance of targeting this pathway to suppress inflammatory responses.

## Acknowledgements

This work was supported by the following NIA grants 5P01AG031862, 5R01AG071861 (PDA), two Glenn Foundation for Medical Research Postdoctoral Fellowship PD19131and AFAR Reboot Fund (N.D.), F32AG066459 and K99AG073450 (KNM).

## Author contributions

N.D and P.D.A. conceptualized and designed the experiments and wrote the manuscript. N.D conducted and led the experiments. N.D, R.A, M.G.T, K.N.M, A.R, A.D, A.H, TL, S.Y, Z.M.C, J.P, M.A, and M.I.R performed the experiments. X.L, C.H.S, K.Y.Y and J.B conducted the computational analysis. V.A and A.R.C performed the proteomics analysis. R.G performed the ZEISS 880 LSM Airyscan confocal Microscopy. D.C.S generated reagents and reviewed the manuscript. S.L.B provided critical input into experimental design, as well as conducted a thorough review and editing of the manuscript.

## Declaration of interests

The authors declare no competing interests.

### Declaration of Generative AI and AI-assisted technologies in the writing process

During the preparation of this work the authors used ChatGPT3.5 in order to enhance readability. After using this tool/service, the authors reviewed and edited the content as needed and take full responsibility for the content of the publication.

## STAR ★ Methods

### Key resources table

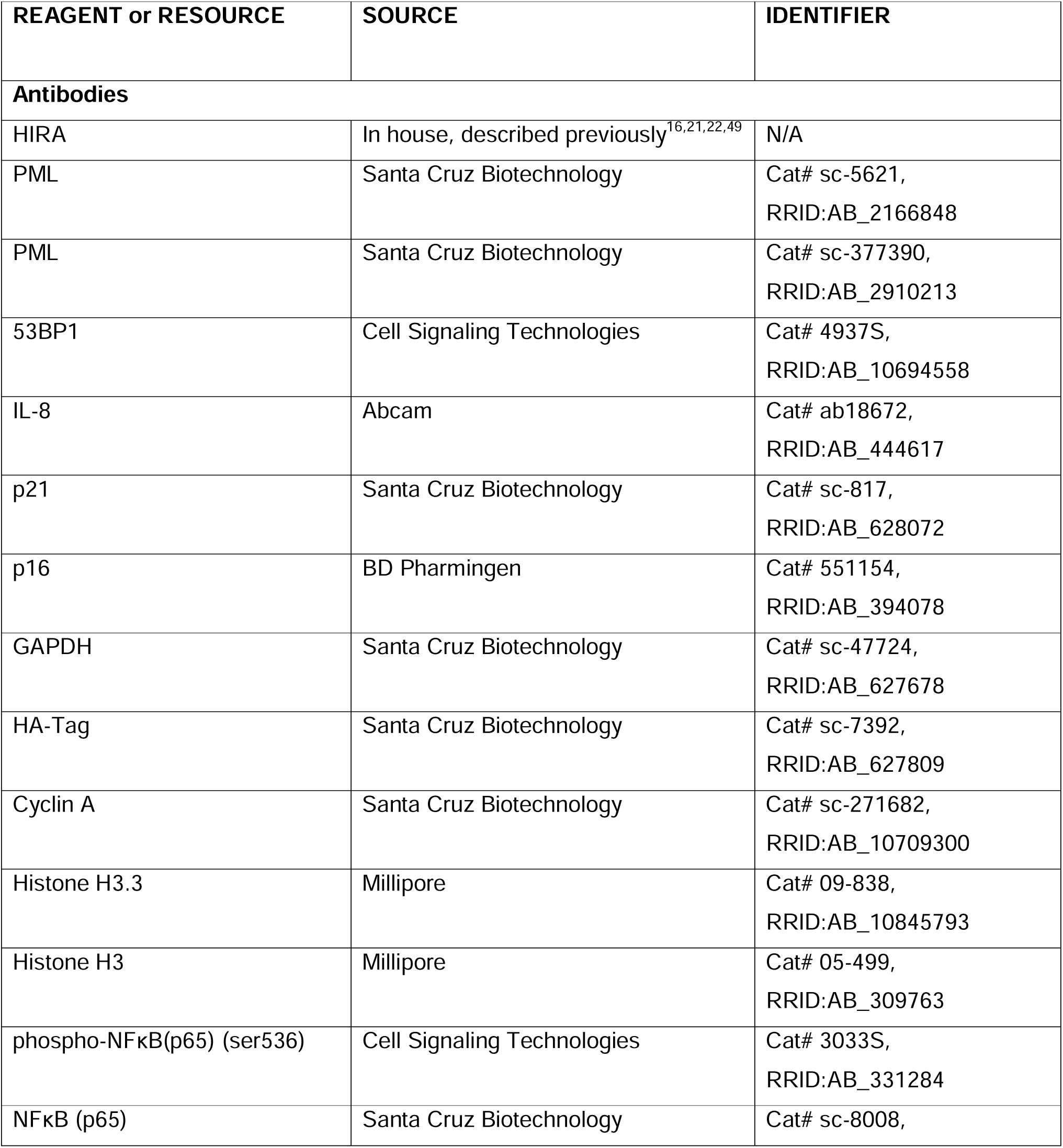

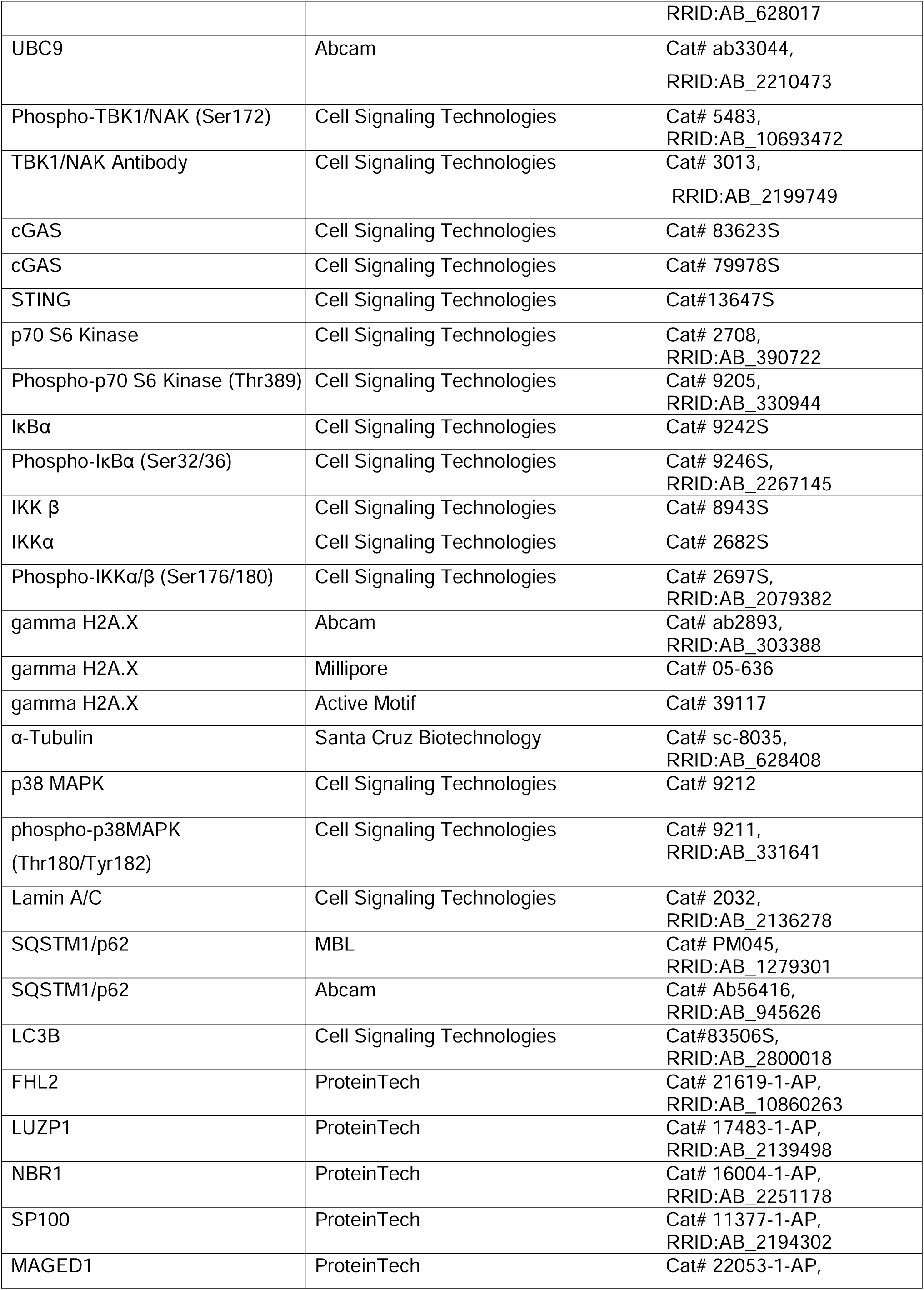

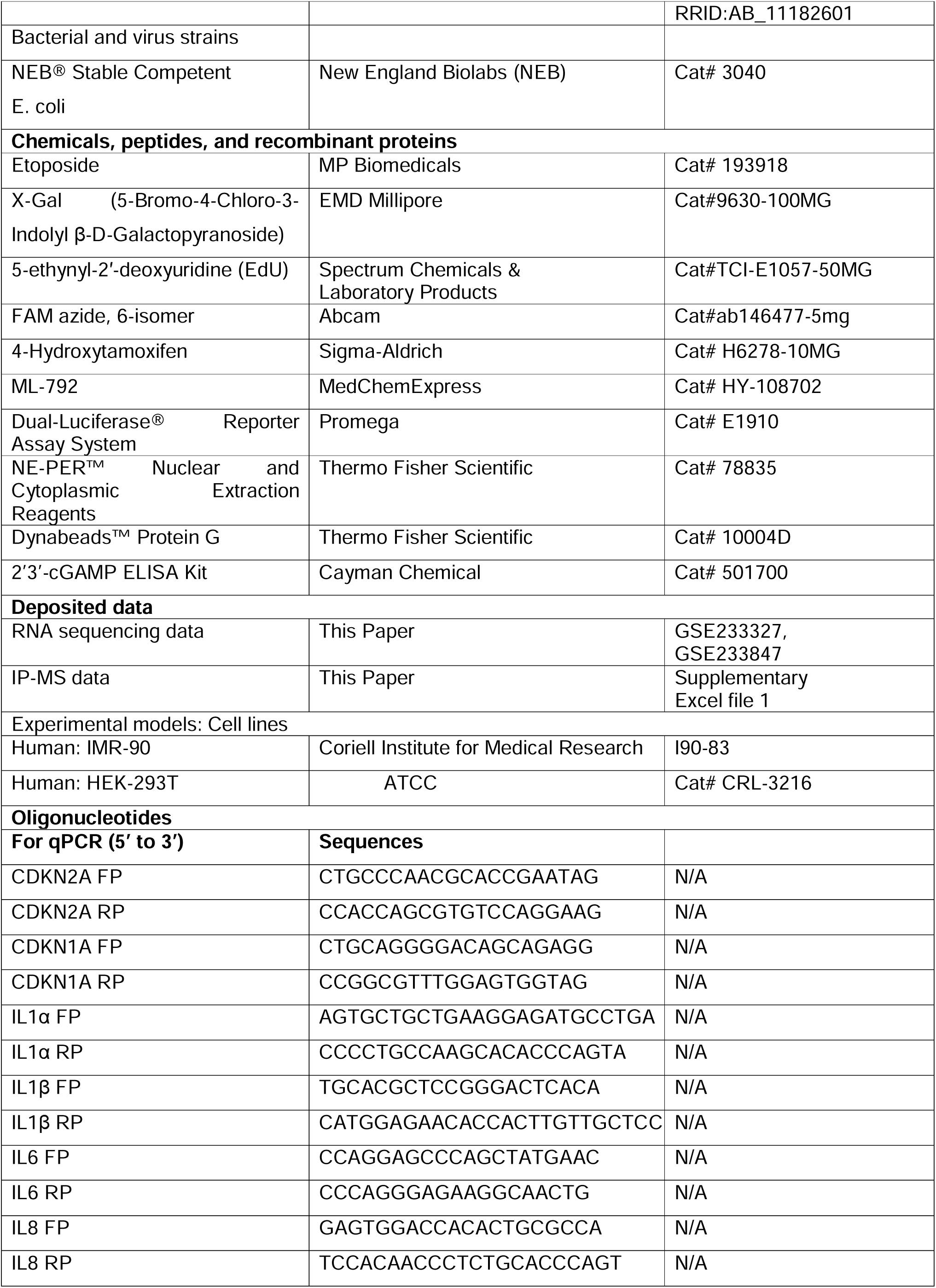

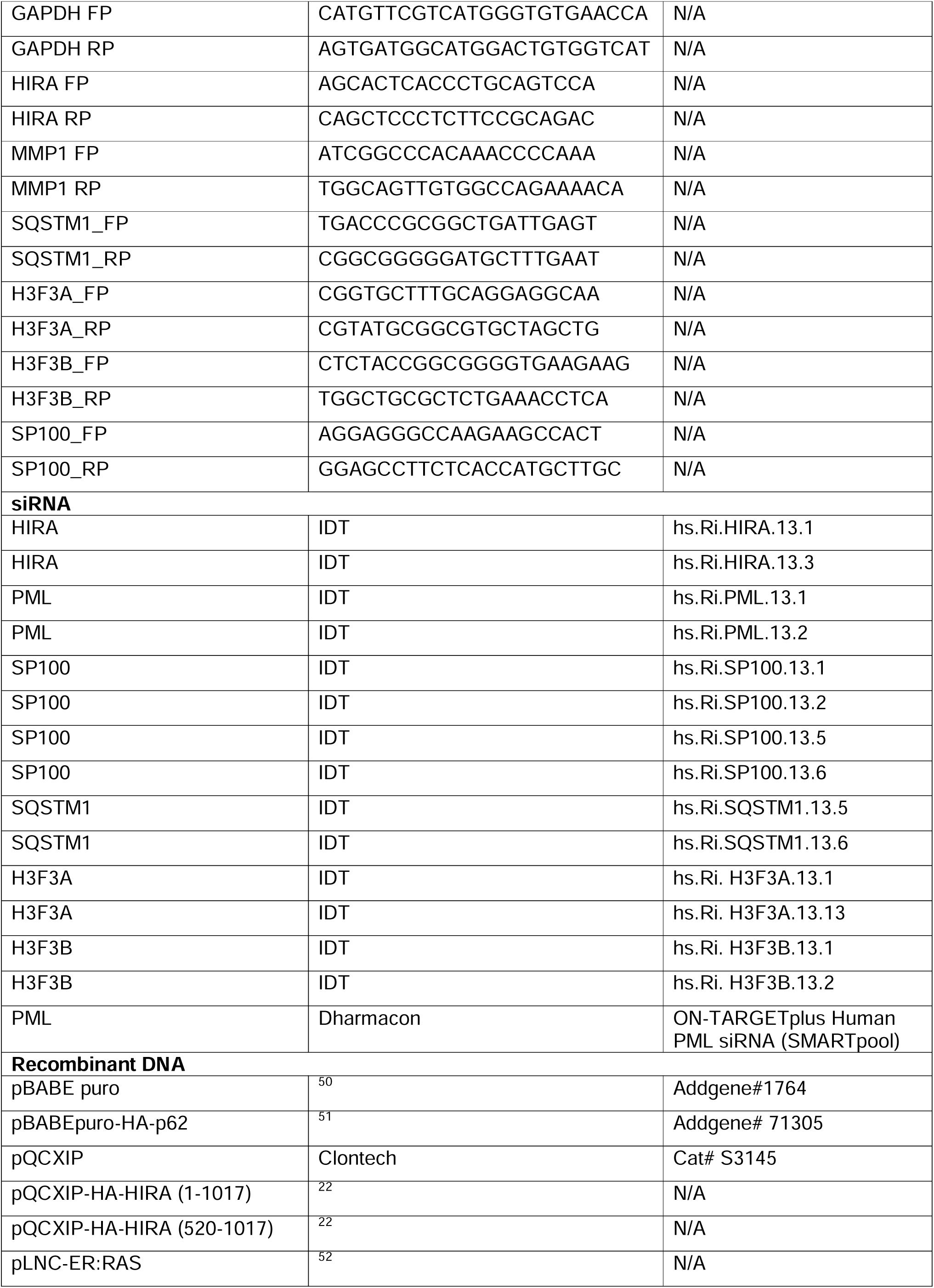

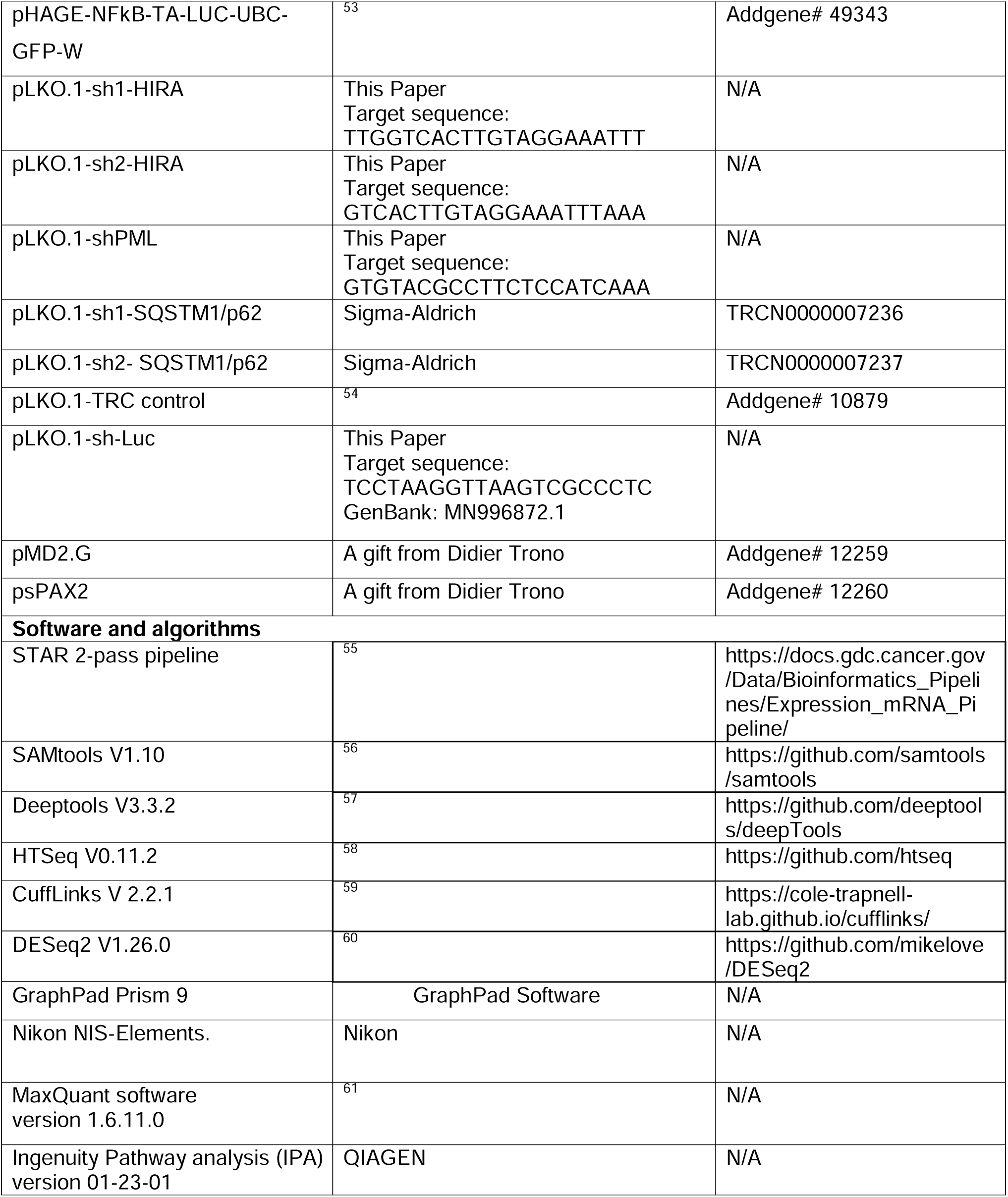

### Resource availability

#### Lead contact

Further information and requests for resources and reagents should be directed to: Prof. Peter Adams

#### Materials availability

All cell lines, DNA constructs and microscopic data are available upon request.

#### Deposited Data

We have deposited all image files and RNA-seq analysis files at Mendeley Database. The link is: https://data.mendeley.com/preview/t896f84hvs?a=46404848-1b10-4203-bf7f-015fded7559d

### Experimental model and study participant details

#### Cell culture and inducing senescence

To culture the cells and induce senescence, primary human IMR-90 fibroblasts were grown in DMEM supplemented with 10% FBS, 100 U/mL penicillin, 100 μg/mL streptomycin, and 2 mM glutamine at 37°C in a humidified atmosphere with 5% CO2 and 3.5% oxygen. To induce senescence, cells were treated with etoposide, a DNA-damaging agent, for 24 hours. For siRNA experiments, etoposide was used at a concentration of 25 μM, while for the rest of the experiments, 50 μM was used. Following this treatment, the etoposide-containing media was replaced with complete media. After seven or eight days, the cells were harvested for senescence assays. Both 25 μM and 50 μM doses, along with the 7–8-day timeline, gave comparable results.

#### Drug treatment in senescent cells

After treating the cells with 50 μM etoposide for 24 hours, the etoposide-containing media was replaced with media containing the drug (ML-792 or LY2090314). The media were then refreshed with new media containing the vehicle and drugs every two days. Seven days post-etoposide treatment, the cells were collected for various assays.

#### Retrovirus Production

Retroviral plasmids was used for the ectopic expression of HA-HIRA^22^, HA-⊿HIRA(520-1017)^22^, and pBABEpuro-HA-p62 (Addgene:71305). Retrovirus production was performed by the SBP Functional Genomics Core Facility using a three-plasmid system consisting of retroviral vector, retro-Gag-Pol, and VSVG, as described earlier^62^. HEK293T cells were co-transfected with the plasmids, and transfected cells were refed with DMEM supplemented with 2.5% FBS. Virus supernatant was collected every 24 hours from the 2^nd^ to 4^th^ day post transfection, and the crude viral supernatant was pooled and filtrated through 0.22-μm membrane. The concentrated viral particles were then purified using 20% sucrose gradient ultracentrifugation at 21,000 rpm for 2 hours, and the resulting pellet was resuspended in 1XPBS, aliquoted, and stored at −80°C. IMR-90 cells were infected with a concentrated viral suspension in the presence of 8 μg/mL polybrene, undergoing two infection cycles for a total duration of 24 hours. The infected cells were then selected using 1 μg/mL puromycin. After 2 days, when the cells reached 80-100% confluency, they were split at a 1:3 or 1:4 ratio based on their confluency level, without puromycin. Once these new plates reached confluency, the cells were again split at a 1:3 or 1:4 ratio as needed. The next day, 50 μM Etoposide was added. After 24 hours of Etoposide treatment, the Etoposide-containing medium was replaced with fresh complete DMEM. Seven- or eight-days post-etoposide treatment, the cells were collected for various assays.

#### Lentivirus Production

The pLKO.1 lentiviral transduction system was used for knocking down HIRA, PML, and SQSTM1/p62. To expresses NF-_K_B luciferase reporter cells, pHAGE NFkB-TA-LUC-UBC-GFP-W lentiviral plasmid (Addgene plasmid # 49343) was used. To produce lentiviral particles, HEK293T cells were transfected with the pLKO.1 constructs and packaging plasmids (pMD2.G, Addgene Plasmid #12259; psPAX2, Addgene Plasmid #12260) using Lipofectamine 2000 (Thermo Fisher Scientific). After six hours, the transfection medium was replaced with fresh culture medium. At 24- and 48-hours post-transfection, the viral supernatants were collected, filtered through a 0.45-μm-pore-size PVDF filter (Millipore), and concentrated by centrifugation at 45,000X g for two hours at 4°C. For infection, IMR-90 cells were incubated with the concentrated viral suspension containing 8 μg/mL polybrene for a duration of 20 hours. The infected cells were subsequently selected using 1 μg/mL puromycin. The infected cells were then selected using 1 μg/mL puromycin. After 2 days, when the cells reached 80-100% confluency, they were split at a 1:3 or 1:4 ratio based on their confluency level, without puromycin. Once these new plates reached confluency, the cells were again split at a 1:3 or 1:4 ratio as needed. The next day, 50 μM Etoposide was added. After 24 hours of Etoposide treatment, the Etoposide-containing medium was replaced with fresh complete DMEM. Seven- or eight-days post-etoposide treatment, the cells were collected for various assays. The shRNA sequences are provided in the supplementary files.

#### si-RNA Transfection

To suppress gene expression in senescent cells, a method involving the transfection of 20 nM Dicer-Substrate Short Interfering RNAs (DsiRNAs) from IDT was employed. This transfection was carried out using Lipofectamine™ RNAiMAX (Invitrogen) following the manufacturer’s protocol. For DsiRNA treatment, we split the cells at a 1:3 or 1:4 ratio based on their confluency. The next day (day 0), we add 25 µM Etoposide. On day 1, we remove the Etoposide-containing medium and transfect the cells with 20 nM DsiRNA using RNAiMax. After an overnight incubation, we replace the DsiRNA medium with fresh complete DMEM (day 2). On day 7, we harvest the cells.

The following DsiRNAs were used for knocking-down: HIRA (hs.Ri.HIRA.13.1, 13.3); PML (hs.Ri.PML.13.1,13.2);SP100(hs.Ri.SQSTM1.13.1,13.2,13.5,13.6);SQSTM1/p62(hs.Ri.SQSTM 1.13.5, 13.6); UBC9(hs.Ri.UBE2I.13.1,13.3); H3.3(pool of H3F3A,H3F3B siRNAs,hs.Ri. H3F3A.13.1,13.13;hs.Ri.H3F3B.13.1,13.2). The ON-TARGETplus Human PML siRNA SMARTPool from Dharmacon was also used for PML knockdown.

#### SA-**β**-Gal assay

The Senescence-associated β-Gal Staining method was adapted from Debacq-Chainiaux et.al^63^. In brief, the cells were fixed at room temperature for 5 minutes in a solution containing 2% formaldehyde and 0.2% glutaraldehyde in PBS. The cells were washed twice with PBS. Subsequently, they were overlaid with a freshly made staining solution and incubated at 37°C for 5-7 hours in a non-CO2 incubator. The staining solution consisted of the following components: 40 mM Na2HPO4 at pH 6, 150 mM NaCl, 2 mM MgCl2, 5 mM K3Fe(CN)6, 5 mM K4Fe(CN)6, and 1 mg/mL X-gal in DMSO. After staining, the cells were rinsed with PBS and then with water to remove salt. The number of β-Gal positive cells was counted under a phase contrast microscope.

#### Edu Assay

The Edu assay was performed based on the method described by Salic and Mitchison^64^. EdU was introduced into the cell media to achieve a final concentration of 10 μM in 24-well plates. Following a 4-hour incubation period, the cells underwent cross-linking with a 10% formalin solution at room temperature for 15 minutes. After aspirating the formalin solution, the cells were subjected to three washes with TBS/0.5% Triton X-100 to permeabilize the cells and quench the formaldehyde. A mixture was prepared, consisting of 2 mL of TBS/0.5% Triton X-100, 2 μL of 50 mM FAM-N3 (6-Carboxyfluorescein Azide) in DMSO, 20 μL of 100 mM CuSO4, and 200 μL of 1.0 M sodium ascorbate. The labeling solution was added to cover the cells adequately, and the mixture was incubated in the dark at room temperature for 30 minutes. Subsequently, the cells were washed three times with TBS/0.5% Triton X-100 and once with PBS. Finally, the cells were stained with DAPI and mounted on slides.

#### Quantitative real-time PCR

The TRIzol method was used to extract total RNA according to the manufacturer’s instructions (Invitrogen). cDNA was prepared from 1 μg of extracted RNA using RevertAidTM reverse transcriptase (Thermo Scientific) according to the manufacturer’s protocol. Quantitative real-time PCR was performed using the PowerTrack™ SYBR Green Master Mix (Applied Biosystems) on the QuantStudioLJ6 Flex Real-Time PCR Systems (Applied Biosystems). The fold change was calculated using the 2^(-ΔΔCt) method on triplicate samples, with GAPDH serving as the housekeeping gene. The primer sequences are provided in the supplementary files.

#### NanoString analysis

Nanostring nCounter analysis was employed to assess SASP gene expression, as previously mentioned^65^. The purified RNA was analyzed on an Agilent 2200 TapeStation using RNA screentape. Subsequently, the samples were run on an nCounter platform using either the nCounter custom human SASP panel, which consisted of 31 canonical SASP genes^2^ and three internal reference genes. The hybridization reactions were carried out following the manufacturer’s instructions from NanoString Technologies. To ensure data quality, field of view counts and binding density measurements were manually extracted from the data and checked against the manufacturer’s protocol to ensure acceptability. Differential expression analysis was conducted using either nSolver to determine log2 fold changes and p-values, which were obtained using Student’s t-test. p-values below 0.05 were considered statistically significant. Heat maps illustrating the normalized counts of specific gene subsets were generated using GraphPad Prism v9.

#### RNA sequencing and analysis

For RNAseq experiments, the RNA extracted using the TRIzol method was further purified using the ammonium acetate precipitation method (Ambion, Catalog: AM9070G). The quality of RNA was assessed using Tapestation from Agilent Technologies. RNA-seq libraries were prepared either by the Genomics Core at SBP Medical Discovery Institute, San Diego or by Novogene, Sacramento, following standard Illumina protocols. RNA-sequencing was conducted as single-read 76 nucleotides in length on the Illumina NextSeq500 platform by the Genomics Core at SBP Medical Discovery Institute (Figures 1 and 2), or on the Illumina NovaSeq PE150 platform (6 G raw data per sample) by Novogene (Figure 3).

Raw fastq files were aligned to hg38 (using STAR 2-pass pipeline^55^). Aligned reads were filtered, sorted and indexed by SAMtools V1.1.0^56^. Genome tracks (bigWig files) were obtained by Deeptools V3.3.2^57^. Raw read counts were obtained by HTSeq V0.11.2 for differential analysis^58^. Differentially expressed genes were obtained by DESeq2 V1.26.0^60^. CuffLinks V2.2.1^59^ was used to compute FPKM values, based on which TPM values were further calculated. TPM values were calculated for gene clustering analysis for both shHIRA/PML and DHIRA experiments. For the clustering of the shHIRA/PML dataset which involved over 15k genes, a two-round hierarchical clustering approach was employed. In the first round, clusters displaying primary patterns were identified, followed by a second round to refine and ensure consistency within each cluster. The clustering of the DHIRA dataset which involved a relatively small set of genes (<1k genes) underwent a single round of hierarchical clustering. (1 - Pearson correlation coefficient) between genes served as the distance measure, and the Ward method was utilized to construct the hierarchical trees, subsequently cut to generate clusters. RNA sequencing data were uploaded to GEO under accession numbers GSE233327 and GSE233847.

#### Immunoblotting

The Western blotting technique was previously described^65^. Cells were lysed in SDS sample buffer (62.5 mM Tris at pH 6.8, 0.01% bromophenol blue, 10% glycerol, and 2% SDS) and incubated immediately at 95°C for 5 minutes. For immunoblotting of STING dimer, cells were lysed in buffer containing 50 mM Tris pH 7.5, 0.5 mM EDTA, 150 mM NaCl, 1% NP40, 1% SDS without any reducing reagent^13^. The proteins were then separated on a 4%-12% Bis-Tris gel (NuPAGE, Thermo Fisher Scientific) through electrophoresis and subsequently transferred to a PVDF membrane. Next, the membranes were blocked in a solution consisting of 5% milk supplemented with 0.1% Tween 20 (TBST) for 1 hour at room temperature. Following the blocking step, the membranes were incubated with primary antibodies overnight at 4°C. The primary antibodies were diluted in 4% BSA in TBST. Afterward, the membranes were washed three times with TBST and incubated with horseradish-peroxidase-conjugated secondary antibodies at room temperature for 1 hour in a solution of 5% milk/TBST. Following another round of washing three times with TBST, the antibody binding was visualized using either SuperSignal™ West Pico or Femto Substrate (Thermo Scientific). All information regarding the antibodies used is provided in the supplementary files.

#### NF-**K**B Luciferase reporter assay

After lentiviral transduction of the lenti-NF-κB-luc-GFP system, IMR-90 cells were further infected with TRC control and two HIRA shRNA-producing pLKO.1 lentivirus particles. Following puromycin selection, the cells were seeded in 24-well plates, and senescence was induced with Etoposide. After seven days, the cells were washed twice with 1X phosphate-buffered saline (PBS) and lysed using 1X passive lysis buffer (Promega). The luciferase activity was measured using the dual luciferase assay kit (Promega) according to the manufacturer’s protocol. Briefly, the lysate was clarified by brief centrifugation, and the firefly luciferase reporter activity in the clear supernatant was measured using a Clariostar Multi-Mode Microplate Reader. GFP fluorescence in the same lysate was measured using the same reader as a normalization control.

#### cGAMP Assay

The 2’3’-cGAMP ELISA Kit (Catalogue no. 501700) from Cayman Chemical was used to quantify 2’3’-cGAMP in the cell lysate following the manufacturer’s protocol. To prepare the cell lysates, Mammalian Protein Extraction Reagent (M-PER™, ThermoFisher Scientific, Catalogue no. 78501) was used. Each sample or standard (50 μL) was added to the appropriate wells of the 2’3’-cGAMP ELISA plate, which was then incubated overnight at 4 LJC with 2’3’-cGAMP-HRP tracer and 2’3’-cGAMP ELISA Polyclonal Antiserum. The plate was washed, and TMB Substrate was added, followed by incubation for 30 minutes at room temperature in the dark. Finally, Stop Solution was added to each well, and the absorbance was measured at 450 nm using a microplate reader.

#### Immunofluorescence Microscopy

Cells were cultured either on cover slips in 24-well plates or in a 96-well plate (PhenoPlate, PerkinElmer) and fixed with 4% paraformaldehyde in PBS for 10 minutes at room temperature. Following fixation, the cells were washed three times with PBS, and the cells were permeabilized with PBS + 0.2% Triton X-100 for 4 minutes at room temperature. Next, the cells underwent blocking with the blocking buffer (PBS + 4% BSA + 1% goat serum) for 1 hour at room temperature, followed by incubation with primary antibodies in blocking buffer for 90 minutes at room temperature. After the incubation, the cells were washed three times with PBS + 1% Triton X-100 and then incubated with secondary antibodies, which were prepared in blocking buffer, for two hours at room temperature. After washing three times with PBS, DAPI (200 ng/mL) in PBS was added for nucleus staining. The cells were incubated for 5 minutes at room temperature in the dark for staining and were subsequently washed three times with PBS. Finally, for coverslips, pre-warmed mounting medium was added to a clean glass slide, and the coverslip with the cells facing down was inverted onto the medium. The edges were sealed with fingernail polish and allowed to dry. The plates or slides were stored at 4°C in a dark environment. In certain nuclear staining experiments, cells were prewashed with EBC lysis buffer (50 mM Tris HCl pH 8.0 + 120mM NaCl + 0.5% NP40 + 5 mM MgCl2 + Protease Inhibitors cocktail) for 30 seconds to reduce the cytoplasmic background before fixation. All information regarding the antibodies used is provided in the supplementary files.

Most of the images were taken automatically in an unbiased manner using a motorized platform by the Nikon Eclipse Ti2 epifluorescence microscope. The image data were processed with Nikon NIS-Elements.

#### Airyscan Super-Res microscopy

High-resolution z-stack images of cells were captured using a ZEISS 880 LSM Airyscan confocal system with an upright stage and a 63x 1.4 NA oil objective. The imaging parameters included a pixel dwell time of 2.05 µs, a pixel size of 20 nm/pixel in SR mode (with a virtual pinhole size of 0.2 AU), and a z-spacing of 0.159 µm. The zoom factor ranged from 2.6 to 3.4. ZEISS Zen software was used to process the images with the Airyscan parameter determined by auto-filter settings. For senescent cells, the Airyscan parameter was increased by 15% compared to the auto-filter settings. DAPI, Alexa Fluor 488, and Alexa Fluor 594 were imaged using the 405 nm, 488 nm, and 561 nm lasers, respectively. Laser power and detector gain were optimized for each staining condition.

#### Cellular fractionation

For cellular fractionation, NE-PER™ Nuclear and Cytoplasmic Extraction Reagents (Catalog: 78835) were used according to manufacturer’s protocol. Briefly, cells pellets were collected as dry as possible by trypsin-EDTA method. To proceed with cytoplasmic and nuclear protein extraction, ice-cold CER I was added to the cell pellet, maintaining the specified reagent volumes. The tube was vortexed vigorously for 15 seconds and then incubated on ice for 10 minutes. Afterward, ice-cold CER II was added to the tube, followed by vortexing for 5 seconds and incubation on ice for 1 minute. The tube was vortexed again for 5 seconds and centrifuged at 16,000 × g in a microcentrifuge for 5 minutes. The supernatant, which contained the cytoplasmic extract. The insoluble fraction containing nuclei was suspended in ice-cold NER and vortexed for 15 seconds. The sample was placed on ice and vortexed every 10 minutes for a total of 40 minutes. Afterward, the tube was centrifuged at 16,000x g in a microcentrifuge for 10 minutes. The resulting supernatant, representing the nuclear extract.

#### Immuno-precipitation

For beads containing HIRA-interacting proteins, a cocktail of mouse monoclonal antibodies (WC15, WC19, WC117, WC119) in approximately equimolar proportions was utilized^16^. For SQSTM1/p62-interacting proteins, an Anti-p62 (SQSTM1) Rabbit polyclonal antibody (MBL, PM045) was used.

First, the cells were scraped from 150-cm2 plates and suspended in 1 mL of EBC500 buffer (50 mM Tris-HCl, pH 8.0; 500 mM NaCl; 0.5% NP40) containing Protease and Phosphatase Inhibitor Cocktail. The resulting lysate was rotated for 30 minutes at 4°C on a rotator and then centrifuged at 12,000x *g* for 10 minutes at 4°C to remove debris and protein concentration of the supernatant was determined using bicinchoninic acid (BCA) protein assay (Thermo Scientific, 23225). For the immunoprecipitation step, Dynabeads™ Protein G (Thermo Scientific, 10004D) was utilized. A volume of 25 μL of Dynabeads per mg of protein was employed, and 2 mL Eppendorf Protein LoBind Tubes were used throughout the procedure. The pooled beads were initially washed twice with 0.5% NP-40 in TBS. Subsequently, the beads were exposed to the antibody for the target protein and isotype control, with a total volume adjusted to 500 μL. The antibody and beads were incubated for 45 minutes at room temperature on a rotator. Following the incubation, the beads were washed three times with 0.5 mL of 0.5% NP-40 in TBS to remove any unbound antibodies. The antibody-bound beads were then added to the lysate and incubated overnight at 4°C on a rotator. Prior to adding the beads, a portion of the lysate was removed to prepare input samples. On a subsequent day, the beads underwent washing either five times with cold NETN buffer (20 mM Tris, pH 8; 1 mM EDTA; 0.5% NP40; 100 mM NaCl) for immunoprecipitation followed by western blot (IP-Western blot), or three times with cold NETN buffer, followed by two times with TBS buffer for immunoprecipitation-mass spectrometry (IP-MS). The DynaMag™-2 Magnet (ThermoScientific,: 12321D) was used for all the washes.

For IP-MS, 10% of the immunoprecipitation solution was collected during the last wash intended for IP-Western blot. The remaining 90% of the beads were collected and subjected to snap-freezing after removing all liquid. Additionally, a western blot was conducted using the 10% aliquot to ensure the quality of the immunoprecipitation prior to mass spectrometry analysis.

#### Immunoprecipitation-mass spectrometry (IP-MS)

Proteomics analysis of immunoprecipitated proteins was performed by the Proteomics Core at SBP Medical Discovery Institute, San Diego. Immunoprecipitated proteins on beads were treated with 8 M urea, 50 mMammonium bicarbonate. Protein disulfide bonds were reduced with 5 mM tris (2-carboxyethyl) phosphine (TCEP) at 30C for 60 min, and cysteines. Subsequently, proteins were alkylated with 15mM iodoacetamide (IAA) in the dark at room temperature for 30 min. Nex, 50mM ammonium bicarbonate was added to dilute ureat to 1M, and proteins were mixed with ammonium bicarbonate and digested with mass spec-grade trypsin/Lys-C mix (1:25 enzyme/substrate ratio) overnight. Following overnight digestion, beads were pulled down, and the digested samples were loaded onto Assa Map reverse-phase resin small pore (C18) cartridges on an Agilent AssayMap Bravo platform. Peptides were eluted with 60% ACN,0.1% FA, and the organic solvent was removed in a SpeedVac concentrator.

Dried peptides were resuspended in 2% ACN, 0.1% FA, and concentration was determined using a NanoDropTM spectrophotometer (ThermoFisher).

The samples were analyzed by LC-MS/MS analysis was performed using a Proxeon EASY-nanoLC system (ThermoFisher) coupled to a Q-Exactive Plus mass spectrometer. Peptides were separated using an analytical C18 Aurora column (75µm x 250 mm, 1.6 µm particles; IonOpticks) at a flow rate of 300nL/min (60C) using 120-min chromatographic gradient: 1% to 5% B in 1min, 6% to 23% B in 72 min, 23% to 34% B in 45 min, and 34% to 48% B in 2 min (A=0.1% FA; B= 80% ACN: 0.1% FA). The mass spectrometer was operated in positive data-dependent acquisition mode. MS1 spectra were measured in the Orbitrap in a mass-to-charge (m/z) of 350-1700 with a resolution of 70,000 at m/z 400. The automatic gain control target was set to 1x10^6 with a maximum injection time of 100ms. Up to 12 MS2 spectra per duty cycle were triggered, fragmented by HCD, and acquired with a resolution of 17,500 and an AGC target of 5x10^4, an isolation window of 1.6m/z, and normalized collision energy of 25. The dynamic exclusion was set to 20 seconds with a 10ppm mass tolerance around the precursor.

#### DDA proteomics analysis

Raw files were processed with MaxQuant software version 1.6.11.0. MS/MS spectra were searched against a Homo sapiens Uniprot protein sequence database (downloaded in January 2020) and GPM cRAP sequences (commonly known protein contaminants). Precursor mass tolerance was set to 20ppm and 4.5ppm for the first search where initial mass recalibration was completed and for the main search, respectively. Product ions were searched with a mass tolerance 0.5 Da. The maximum precursor ion charge state used for searching was 7. Carbamidomethylation of cysteine was searched as a fixed modification, while oxidation of methionine and acetylation of protein N-terminal were searched as variable modifications. Enzyme was set to trypsin in a specific mode and a maximum of two missed cleavages was allowed for searching. The target-decoy-based false discovery rate (FDR) filter for spectrum and protein identification was set to 1%.

#### Statistics

The statistical significance was analyzed utilizing GraphPad Prism 9 Software. In most cases, the value for the proliferation control was considered as 1, following convention, and the fold change of senescent cells was calculated. To determine whether the fold change significantly differed from 1, the one-sample t-test was employed. When comparing two datasets, unpaired two-tailed Student’s t-test was performed. For the comparison of more than two datasets, the one-way ANOVA test was used. A significance level of p<0.05 was considered to determine statistical significance. The data are expressed as Mean±SD from three biological replicates and are representative of at least two independent experiments. The symbols #,IZ, IZIZ, IZIZIZ, IZIZIZIZ indicate significance levels of p<0.01, p<0.05, p<0.01, p<0.001,and p<0.0001, respectively.

