## Supplementary figures and images for "Histone chaperone HIRA, Promyelocytic Leukemia (PML) protein and p62/SQSTM1 coordinate to regulate inflammation during cell senescence"

### S1

#### Figure S1:HIRA accumulates in PML bodies in senescent cells.

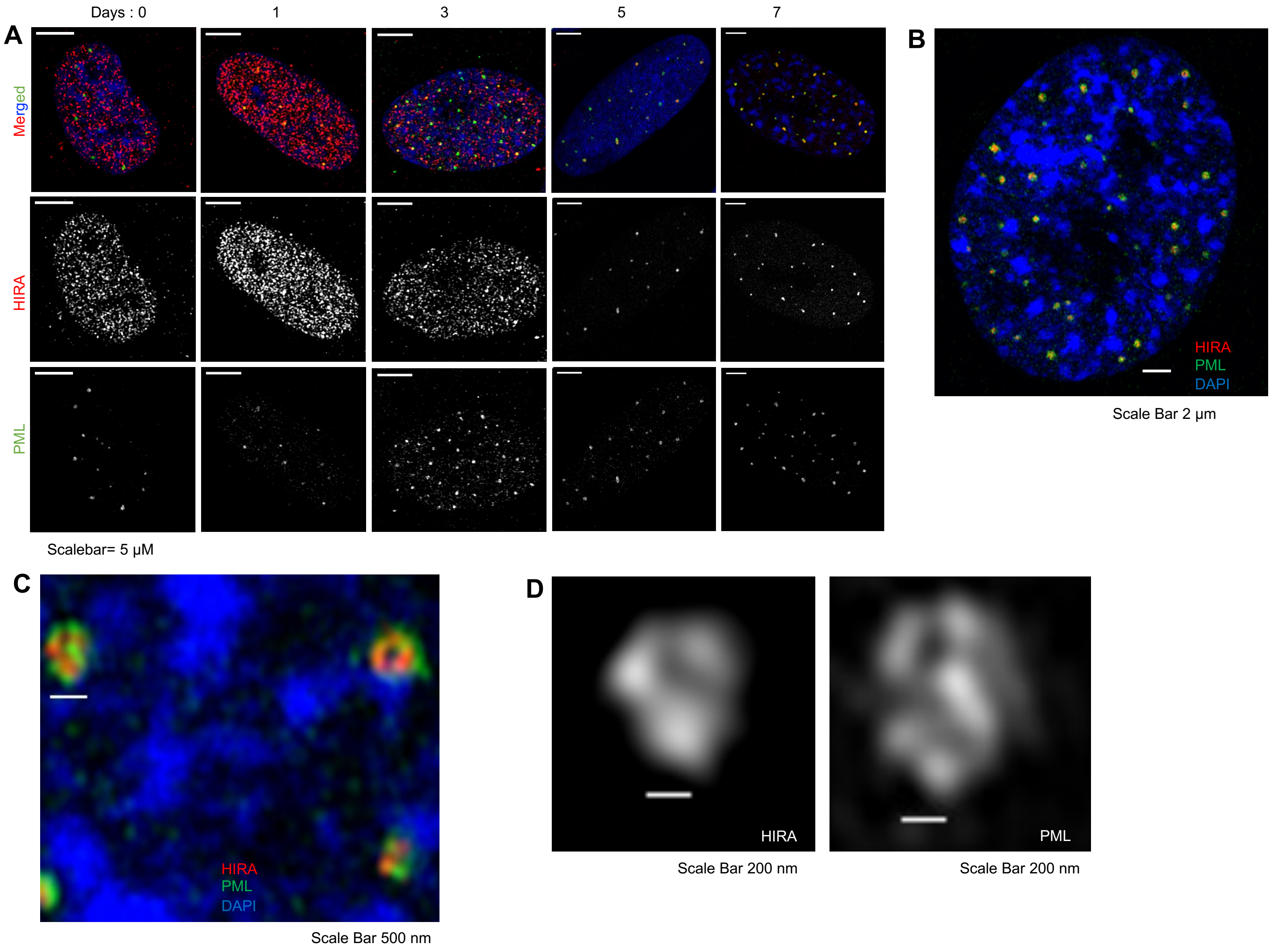

### S3

##### Figure S3: PML and HIRA foci are frequently wrapped by DNA-SCARS.

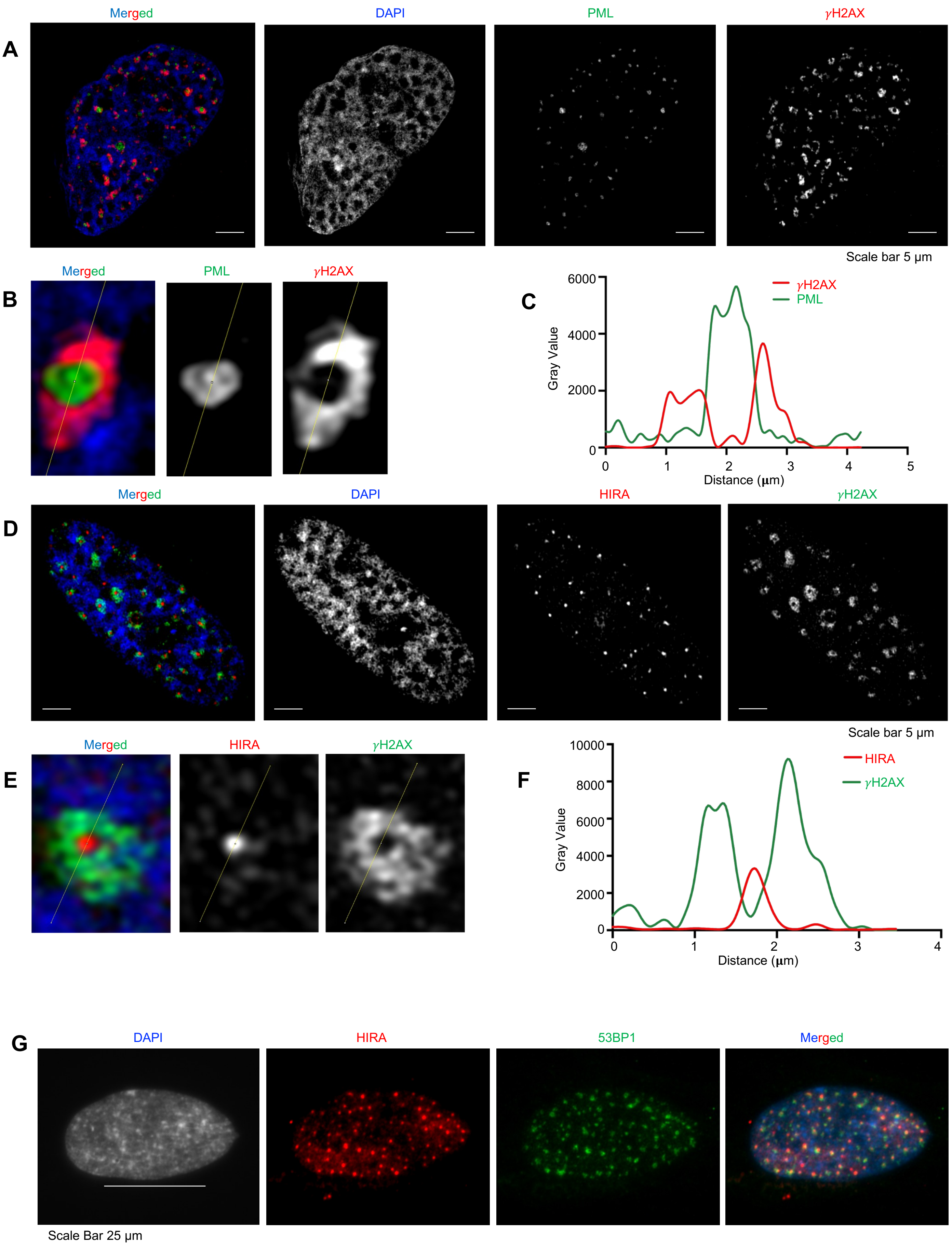

### S7

**Figure S7: cGAS expression in senescent cells is negatively associated with p62 Expression.**

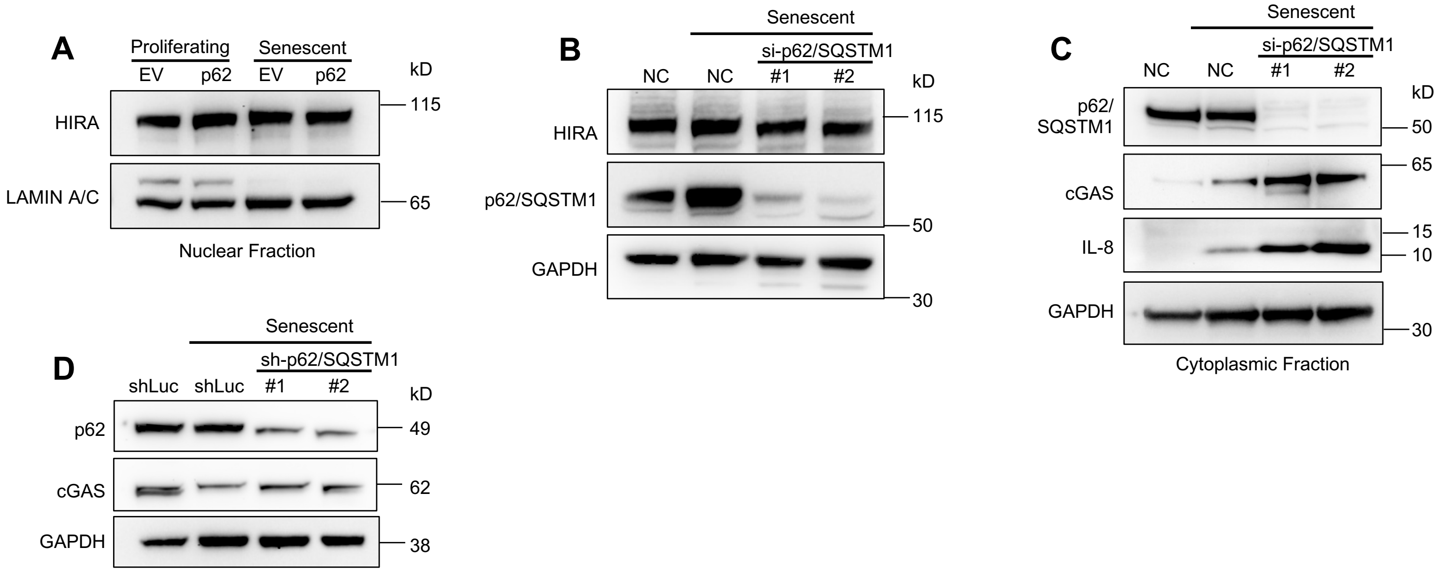

### S8

##### Figure S8: HIRA exhibits a physical interaction with the autophagy regulator p62.

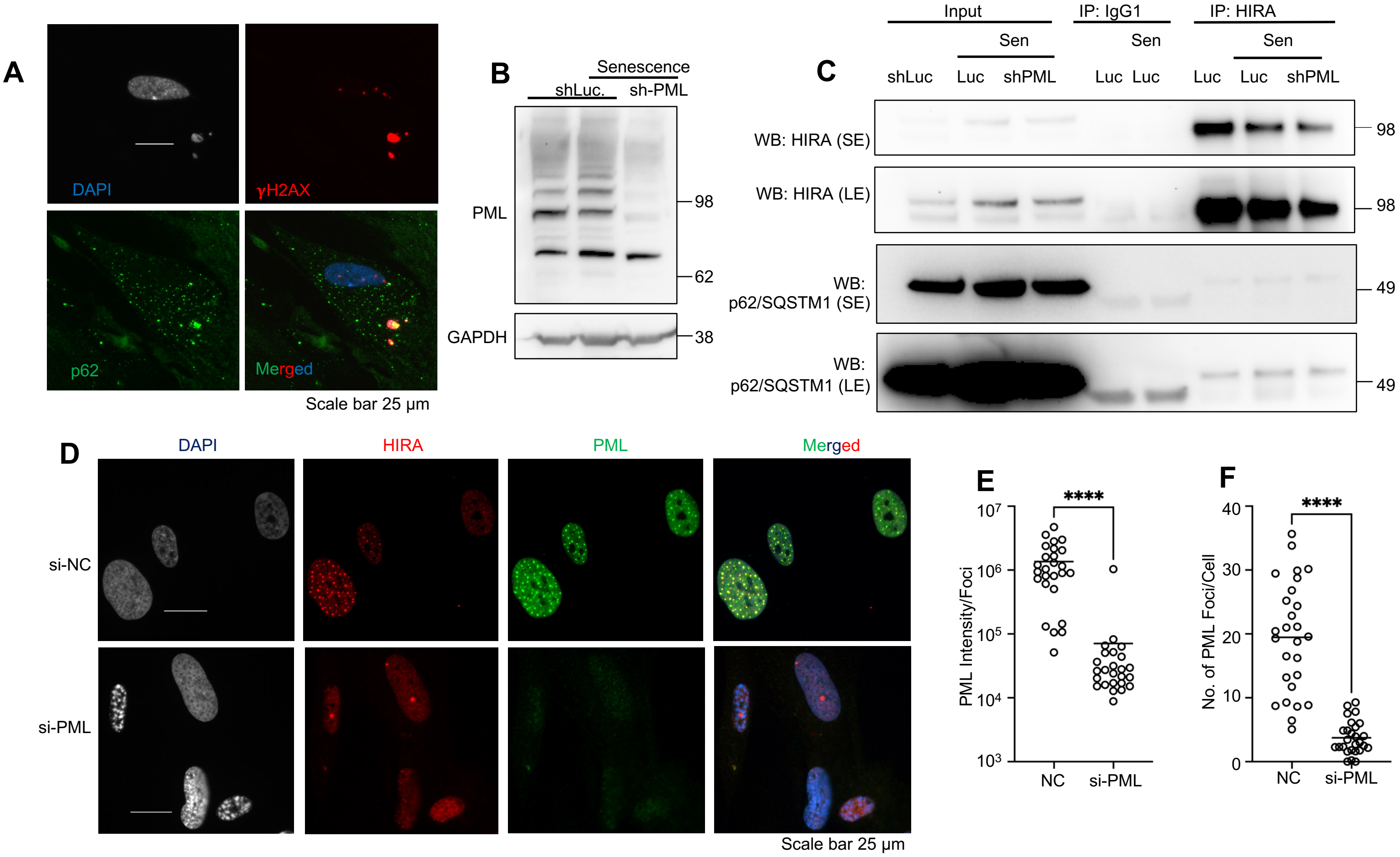
