## Supplementary material for "Histone chaperone HIRA, Promyelocytic Leukemia (PML) protein and p62/SQSTM1 coordinate to regulate inflammation during cell senescence": S2

**Figure S2: PML, HP1α, and SP100 exhibited high resistance to dissolution in 1,6-hexanediol treatment, whereas HIRA exhibited very low resistance.**

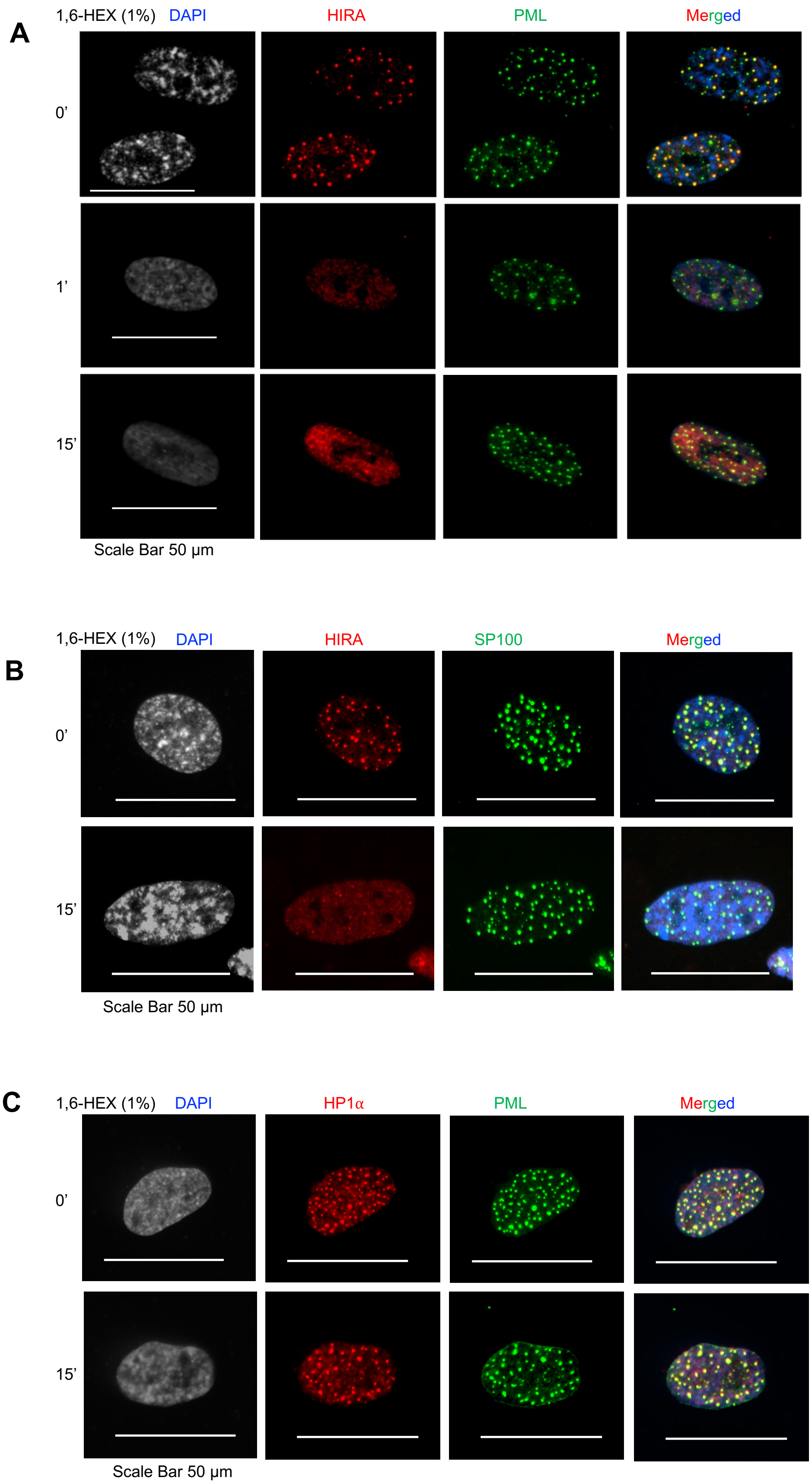
