## Supplementary material for "Histone chaperone HIRA, Promyelocytic Leukemia (PML) protein and p62/SQSTM1 coordinate to regulate inflammation during cell senescence": S4

**Figure S4: HIRA and PML are necessary for SASP expression but not for proliferation arrest in different senescence models**

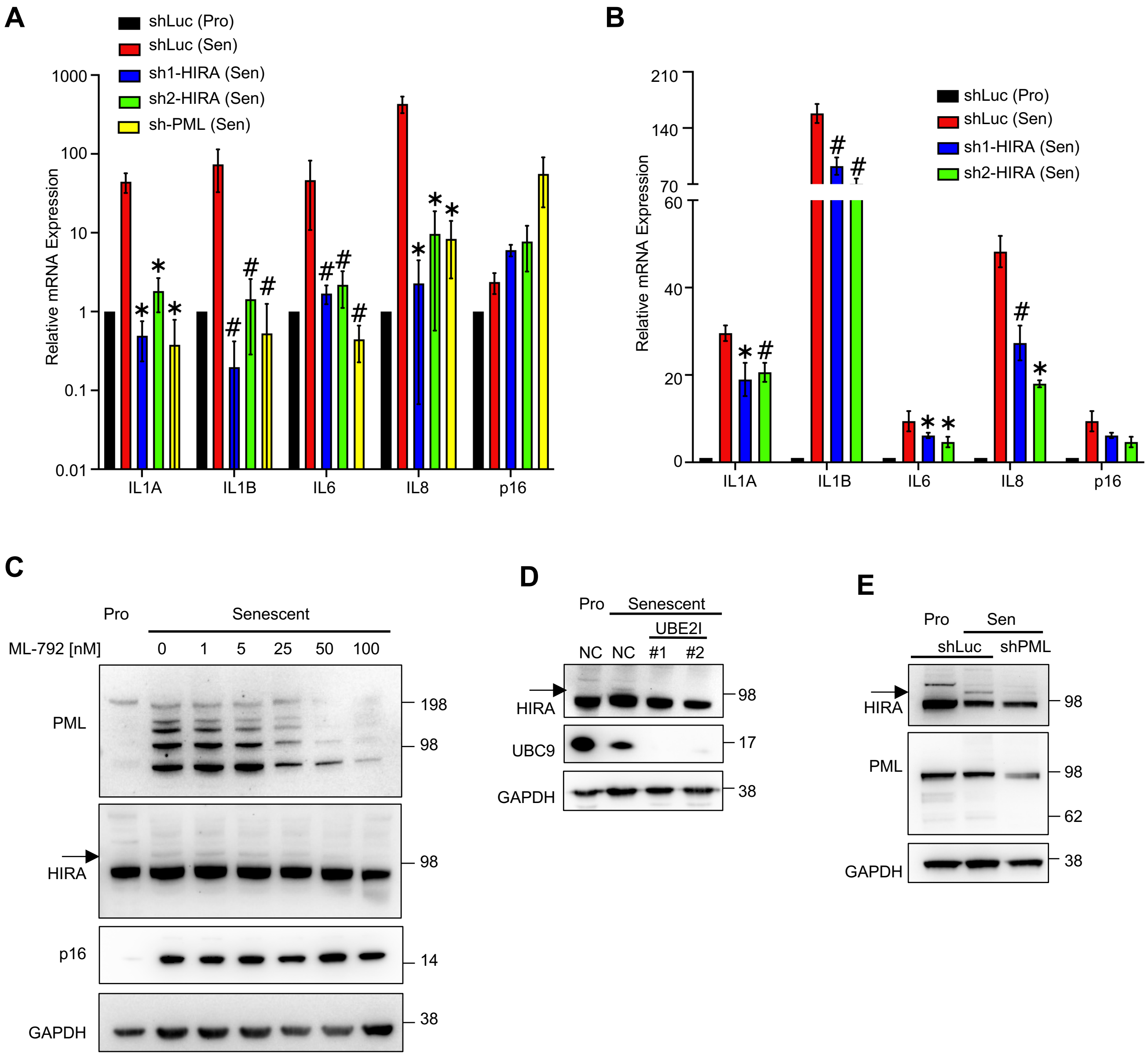
