## Supplementary material for "Histone chaperone HIRA, Promyelocytic Leukemia (PML) protein and p62/SQSTM1 coordinate to regulate inflammation during cell senescence": S5

### Figure S5: More than 1/3 of the genes affected by HIRA and PML shRNA were also altered by ΔHIRA.

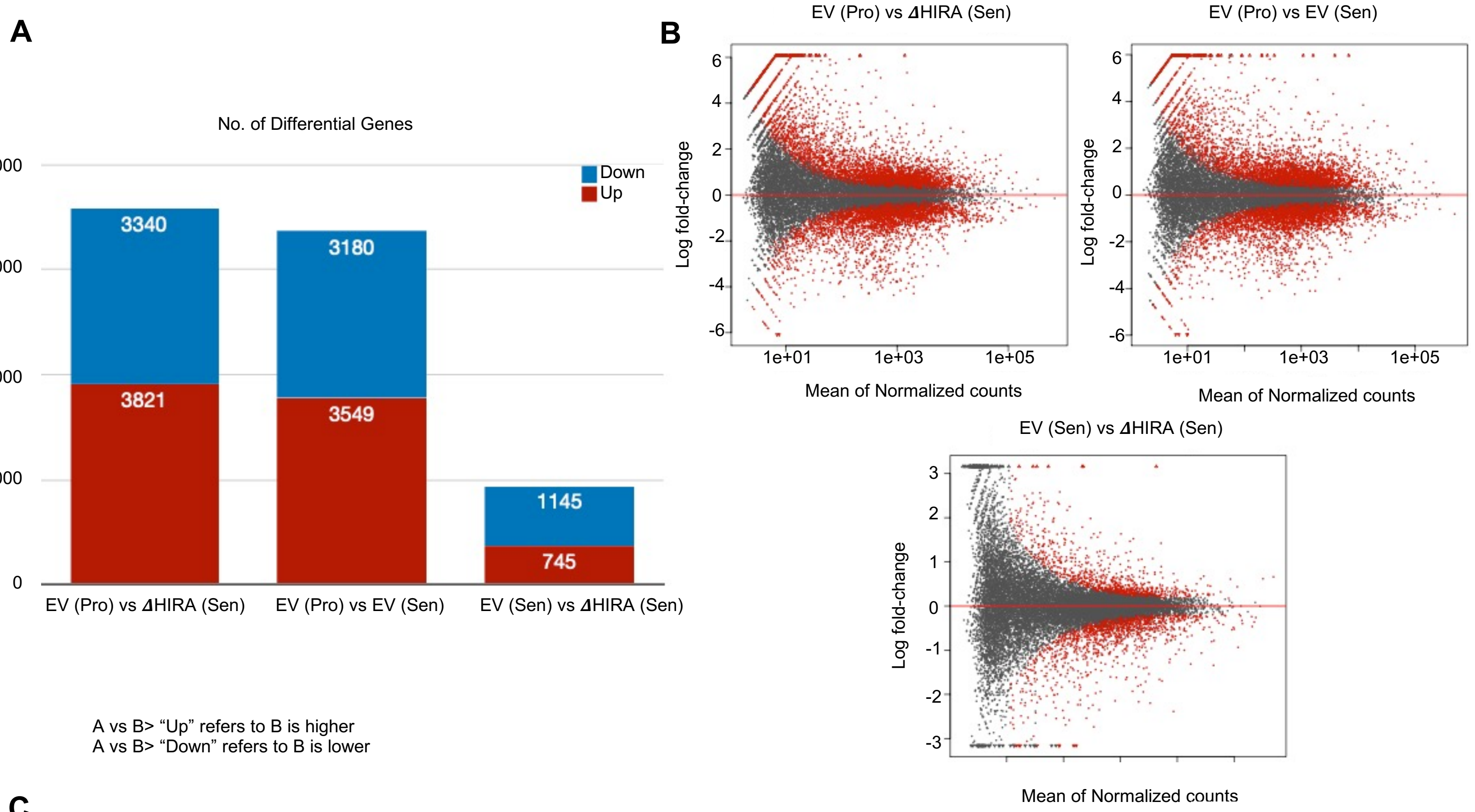


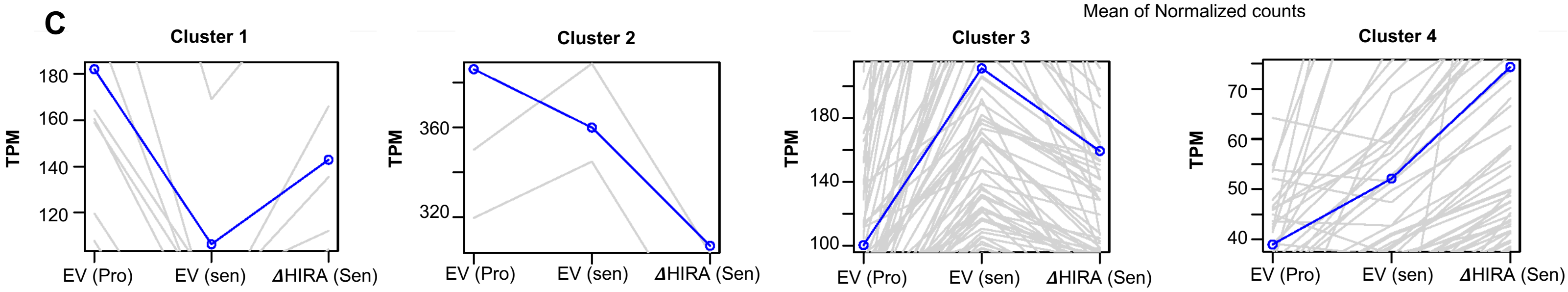


| Ingenuity Canonical Pathways | -Log (p-value) |
|---|---|
| Tryptophan catabolism | 3.76 |
| Mevalonate Pathway I | 3.62 |
| Transport of inorganic cations/anions and amino acids/ oligopeptides | 3.17 |
| Superpathway of Cholesterol Biosynthesis | 2.98 |
| RHOA Signaling | 2.95 |
| Fatty Acid β-oxidation I | 2.84 |
| Collagen biosynthesis and modifying enzymes | 2.26 |
| Remodeling of Epithelial Adherens Junctions | 2.25 |
| IL-17A Signaling in Fibroblasts | 2.11 |
| Superpathway of Serine and Glycine Biosynthesis I | 2.09 |

| Ingenuity Canonical Pathways | -Log (p-value) |
|---|---|
| Eukaryotic Translation Initiation | 23.2 |
| Eukaryotic Translation Termination | 21.9 |
| Eukaryotic Translation Elongation | 21.8 |
| EIF2 Signaling | 17.8 |
| Major pathway of rRNA processing in the nucleolus and cytosol | 16.7 |
| mTOR Signaling | 5.1 |
| Regulation of eIF4 and p70S6K Signaling | 4.57 |
| Interferon alpha/beta signaling | 3.45 |
| Interferon Signaling | 3.19 |
| Purine Nucleotides De Novo Biosynthesis II | 2.93 |

| Ingenuity Canonical Pathways | -Log (p-value) |
|---|---|
| Interleukin-10 signaling | 16.2 |
| Interleukin-4 and Interleukin-13 signaling | 12.7 |
| IL-17A Signaling in Fibroblasts | 11.8 |
| Wound Healing Signaling Pathway | 10.1 |
| Hepatic Fibrosis Signaling Pathway | 10 |
| Pathogen Induced Cytokine Storm Signaling Pathway | 9.95 |
| Role of Macrophages, Fibroblasts and Endothelial Cells in Rheumatoid Arthritis | 8.3 |
| Role of Pattern Recognition Receptors in Recognition of Bacteria and Viruses | 8.19 |
| Interferon alpha/beta signaling | 8.11 |
| --- | --- |
| CGAS-STING Signaling | 6.39 |
| HMGB1 Signaling | 6.19 |
| IL-6 Signaling | 6.01 |

| Ingenuity Canonical Pathways | -Log (p-value) |
|---|---|
| Ion channel transport | 5.88 |
| Extracellular matrix organization | 4.2 |
| VDR/RXR Activation | 4.04 |
| RHO GTPase cycle | 4.02 |
| Cholesterol Biosynthesis I | 3.67 |
| Cholesterol Biosynthesis II | 3.67 |
| Cholesterol Biosynthesis III | 3.67 |
| Cholesterol biosynthesis | 2.8 |
| CGAS-STING Signaling | 2.68 |
| Iron uptake and transport | 2.67 |
| Post-translational protein phosphorylation | 2.47 |
| Collagen biosynthesis and modifying enzymes | 2.44 |
| Collagen biosynthesis and modifying enzymes | 2.44 |
| Vitamin D metabolism | 2.27 |
| Inhibition of Matrix Metalloproteases | 2.24 |
| Regulation of Insulin-like Growth Factor (IGF) transport and uptake by IGFBPs | 2.2 |
