## Supplementary material for "Histone chaperone HIRA, Promyelocytic Leukemia (PML) protein and p62/SQSTM1 coordinate to regulate inflammation during cell senescence": S6

**Figure S6: HIRA and PML are not necessary for the regulation of certain effectors involved in SASP.**

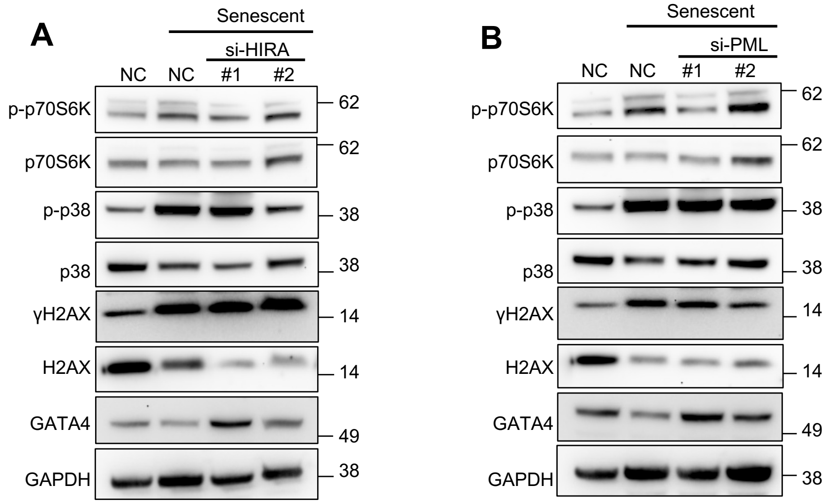
